# Membrane damage during *Candida albicans* epithelial invasion is localized to distinct host subcellular niches

**DOI:** 10.1101/2025.05.13.653632

**Authors:** Laura Marthe, Noa Conan, Anouck Shekoory, Patricia Latour-Lambert, Jean-François Franetich, Catherine Pioche-Durieu, Rémi Le Borgne, Jean-Marc Verbavatz, Martin Larsen, Allon Weiner

## Abstract

*Candida albicans* is an opportunistic fungal pathogen normally found as a commensal in human mucosa. In susceptible patients, its hyphal filamentous form can invade and damage host epithelium leading to local or systemic infection. The secreted fungal toxin candidalysin is required for epithelial damage, although both hyphal extension and candidalysin are required for inducing maximal infection damage, for reasons that are not well understood. Here we study the interplay between hyphal extension and epithelial damage *in vitro* using live cell imaging combined with damage-sensitive reporters and volume electron microscopy. We first quantify candidalysin-induced host membrane rupture and host cell death at the single cell level and determine the three-dimensional architecture of invasion in our experimental system. We then show that the spatial organization of host membrane rupture is regulated by candidalysin and is predictive of host cell death. Finally, we demonstrate that membrane rupture occurs at two distinct host subcellular niches: the juxtanuclear region and the cell-cell boundary, with the latter the main driver of subsequent host cell death. Based on these results, we propose that *C. albicans* invasion damages epithelial membranes in a sequential process in which candidalysin secretion first acts to weaken host membranes enveloping invading hyphae, followed by membrane rupture occurring as hyphae traverse two distinct host subcellular niches.

## Introduction

The oral, genital and intestinal mucosa of most healthy individuals are colonized by the fungal pathobiont *Candida albicans* as part of the normal commensal microflora[1]. *C. albicans* causes superficial infections such as oral or vaginal thrush, and in susceptible hosts can invade the intestinal epithelial barrier and enter the bloodstream, leading to severe systemic infection[2]. *C. albicans* transforms from an ovoid yeast cell to a filamentous hyphal form in response to various environmental cues, a transition that facilitates tissue invasion and subsequent host damage[3].

The first barrier encountered by *C. albicans* during infection is the host epithelium. After attachment of the yeast form to the epithelial surface, hyphal growth is initiated, followed by invasion into the epithelial layer, mainly via an intracellular (transcellular) route[4,5]. As hyphae extend, they are tightly enveloped by the plasma membrane of the invaded host cell, forming a structure known as the ‘invasion pocket’[6,7]. During invasion into Caco-2 cells, a model for enterocytes, extending hyphae can expand the invasion pocket into neighboring host cells without membrane breaching, forming a multi-layered structure termed *Candida albicans* trans-cellular tunnel (CaTCT)[5]. Hyphal extension within the invasion pocket or trans-cellular tunneling ultimately results in host membrane damage, followed by host cell death and epithelial layer degradation[4]. Hyphal extension *per se* is thought to cause little or no host damage despite the extensive remodeling of invasion pocket membranes during invasion[6,8]. Instead, damage is dependent on candidalysin, a secreted fungal peptide toxin encoded by the gene *ECE1*, that targets host glycosaminoglycans and induces the formation of pores within host membranes[6,9,10]. *C. albicans* strains that do not express candidalysin adhere and invade normally, but are unable to damage host cells[6,7]. As the exogenous addition of synthetic candidalysin peptide directly to host cells results in significantly less damage compared to that observed during *C. albicans* hyphal invasion (unless extremely high peptide concentrations are used), it is thought that both candidalysin secretion and hyphal invasion are required for *C. albicans* to induce its full damage potential[6,7,11]. Mogavero et al. have shown using anti-candidalysin llama V_H_Hs (nanobodies) that during infection of oral epithelial cells, candidalysin is secreted within the invasion pocket[7]. They go on to propose that candidalysin accumulation within the invasion pocket allows the toxin to reach a concentration sufficient for host membrane damage, thus explaining the observed difference in damage between exogenously added candidalysin and hyphal invasion. Additionally, Moyes et al. have reported that different concentrations of exogenously added candidalysin induce a differential host response in epithelial cells: signal transduction through MAPK, p38/MKP-1 and c-Fos, resulting in the production of immune cytokines by sub-lytic candidalysin concentrations, and a calcium influx resulting in the release of damage associated cytokines by lytic candidalysin concentrations[6]. Thus, it was proposed that candidalysin has dual functionality in a concentration-dependent manner, and that during infection, it is candidalysin accumulation within the invasion pocket that drives a matching dual innate immune response: pathogen recognition before appreciable host membrane damage, followed by a host response to membrane damage[6,12].

Current understanding of *C. albicans* epithelial invasion and damage is based mostly on end-point microscopy following sample fixation as well as population-level damage assays such as LDH release[4,11,13]. While highly informative, such approaches are limited in their ability to assess the dynamics and precise spatial organization of these processes. In this work we explore the interplay between epithelial invasion, damage and candidalysin secretion using an *in vitro* epithelial infection model, live cell imaging in combination with real-time damage-sensitive reporters and volume electron microscopy[5]. Our approach allows for direct, single-cell level quantitative investigation of *C. albicans* epithelial infection at different spatial and temporal scales. We begin by analysis of the global dynamics of host membrane rupture and cell death using a *C. albicans* wild-type strain and a mutant strain that does not express candidalysin. We go on to describe the three-dimensional architecture of invasion and the spatial organization of host membrane rupture. Finally, we demonstrate that membrane rupture occurs at two distinct host subcellular niches, each associated with specific structural and functional attributes. Based on these results, we propose a new ‘sequential’ mechanism for the infliction of damage by *C. albicans* during epithelial infection and discuss how it relates to existing concepts in the field.

## Results

### Experimental pipeline for studying *C. albicans* invasion and damage using live cell imaging combined with damage-sensitive reporters

In order to track the dynamics and spatial organization of *C. albicans* invasion and damage at the single-cell level, we established an *in vitro* experimental epithelial infection model designed for quantitative live cell imaging in three-dimensions in combination with damage-sensitive reporters (**Fig 1A, 1B**). TR146 oral epithelial cells, widely used to study *C. albicans* infection[6,9,14,15], were engineered to express the membrane rupture marker Galectin-3 linked to the fluorescent protein mOrange[5,16,17]. Galectin-3 is a small soluble protein localized to the cytosol, which binds β-galactose-containing carbohydrates typically found on the luminal side of intracellular vesicles and at the outer leaflet of the plasma membrane. By tracking mOrange-Galectin-3 (hence referred to as Gal-3), the location and spatial organization of membrane rupture can be identified in real-time, as once a host membrane is breached, cytosolic Gal-3 flows through the rupture site and onto the cell surface where it remains bound. This approach enables the detection of significant membrane damage generally considered ‘catastrophic’, but is incapable of detecting smaller, ‘non-catastrophic’ rupture such as those induced by bacterial toxins[18,19]. Following seeding in a microplate (96-well plate), confluent epithelial cells are stained with CellMask Deep Red (CM), which labels the host plasma membrane and endocytic compartment, including *C. albicans* induced ‘invasion pockets’, but not *C. albicans* membranes[5]. TR146 cells stained by CM present clear cell-cell boundaries (CCBs) and a subcellular region rich in vesicles adjacent to the nucleus we refer to as ‘the juxtanuclear region’ (JNR). Host cells are also stained with SYTOX green (SYTOX), a nucleic acid stain that is impermeant to live cells but marks the nucleus of lysed dead cells. In all experiments, two *C. albicans* strains are studied in parallel (in different wells): the wild-type (WT) prototroph BWP17 and *ece1*Δ/Δ, a mutant strain that does not express the fungal toxin candidalysin [6,11,20]. Epithelial cells are infected at a ‘multiplicity of infection’ (MOI) of 0.001, whereby a single *C. albicans* germ cell, hence referred to as an ‘inoculating yeast’ (IY), is introduced for every thousand epithelial cells. Live cell imaging is then launched for 12 hours with 10 min imaging intervals (40X objective) or 24 hours with 30 min imaging intervals (20X objective), depending on the specific experiment, followed by image analysis and quantification. Imaging is performed in parallel at multiple infection sites within each well and in multiple wells, allowing for direct comparison between different conditions, such as WT and *ece1*Δ/Δ strains or uninfected controls. Images are acquired in four channels: Brightfield (BF), SYTOX, Gal-3 and CM (**Fig 1B**). Individual membrane rupture events are detected as transient local increases in Gal-3 signal (hence ‘Gal-3 recruitments’) along the trajectories of extending hyphae detected in the CM channel (**Fig. 1B, green inset**).

**Fig. 1.**
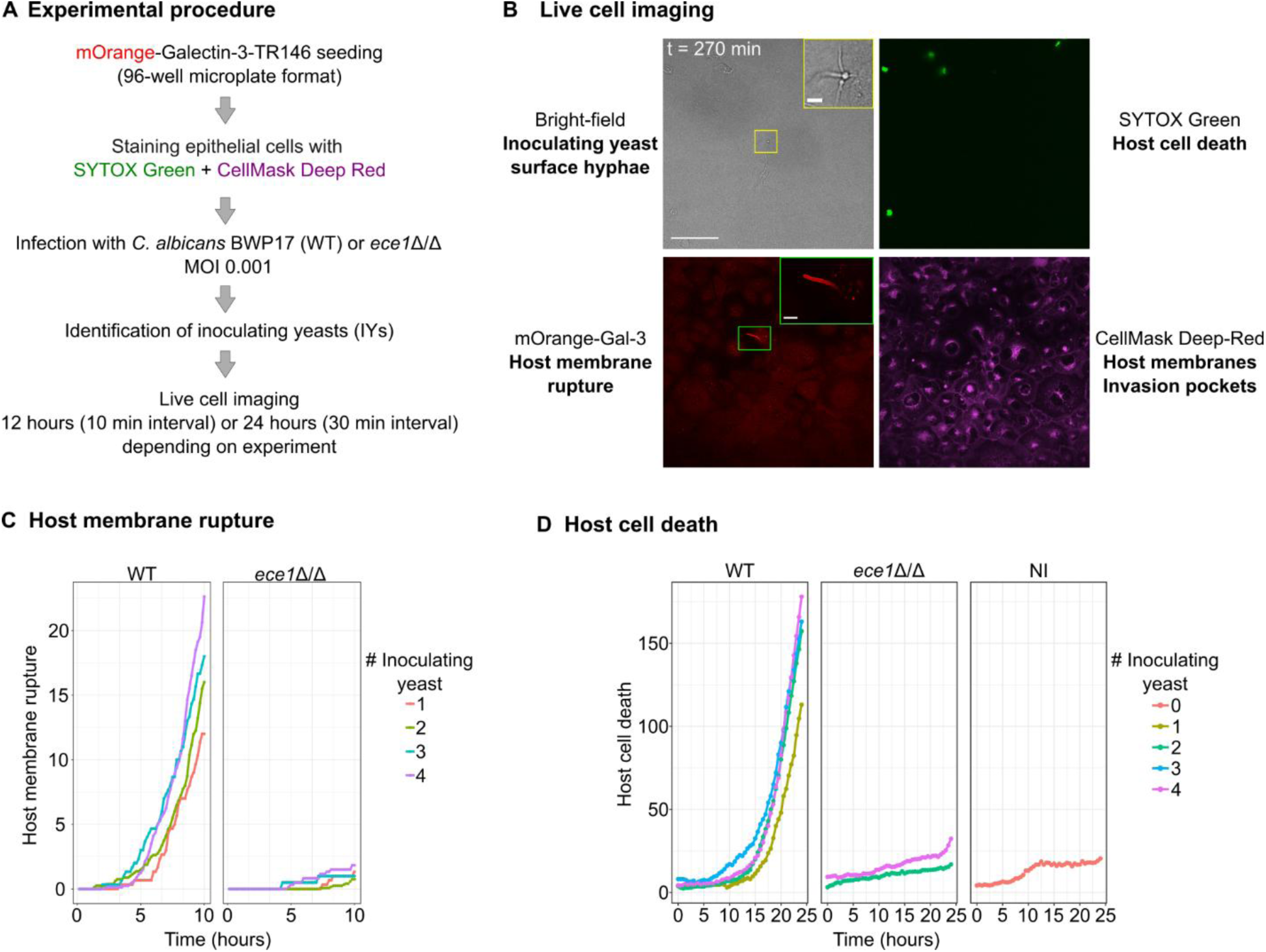
Live cell imaging combined with damage-sensitive reporters used to study host membrane rupture and host cell death during *C. albicans* infection of TR146 oral epithelial cells. (A) Experimental procedure. Confluent TR146 cells expressing mOrange-Galectin-3 plated in a 96-well microplate are stained with the host membrane marker CellMask Deep Red and the cell death marker SYTOX green, followed by the addition of *C. albicans* wildtype and *ece1*Δ/Δ strains at a MOI of 0.001. Inoculating yeast (IY) identification is followed by live cell imaging for either 12 hours with 10 min intervals or 24 hours with 30 min intervals, depending on the experiment. (B) Live cell imaging data is acquired in four channels: brightfield channel - recording of IY and hyphal surface extension, inset shows magnified view of IY and hyphae; SYTOX green - host cell death; mOrange-Gal-3-host membrane rupture as indicated by galectin-3 recruitments, inset shows magnified view of membrane rupture event; CellMask Deep-Red - host membrane organization including invasion pockets induced by *C. albicans* hyphae. *C. albicans* membranes are not stained by CellMask in our experimental procedure. (C) Total host membrane rupture over time as indicated by galectin-3 recruitments using WT strain and *ece1*Δ/Δ strain that does not express candidalysin. Individual Gal-3 recruitments were recorded every 10 minutes during the first 10 hours of infection. A total of 60 WT IY linked to 496 Gal-3 recruitments and 57 *ece1*Δ/Δ IY linked to 33 Gal-3 recruitments from 4 independent experiments were recorded. For each IY number (1-4), the average number of Gal-3 recruitments at each timepoint is presented (D) Total host cell deaths over time as indicated by SYTOX green labeling using WT and *ece1*Δ/Δ strains. Non-infected (NI) wells are used as a control (IY = 0). Individual SYTOX green labeling events were recorded every 30 min during 24 hours. A total of 47 WT IY linked to 3107 SYTOX green labeling events, 46 *ece1*Δ/Δ IY linked to 376 SYTOX green labeling events and 224 SYTOX green labeling events in non-infected wells from 3 independent experiments were recorded. For each IY number (1-4), the average number of SYTOX green labeling events at each timepoint is presented. Scale bars are 100 µm and 10 µm in insets. See also **S1 movie and S2 movie.**

A unique feature of our experimental design is the ability to track epithelial infection by a low number of IYs over time within a well-defined ‘infection site’ (**Fig. 1B, yellow inset, S1 Fig**). Before the launching of a live cell imaging experiment (t = 0), a small group of IYs (1-4) clumped together are manually identified and centered to the middle of the microscope field-of-view (FOV) so that no additional IYs are observed. Thus, all hyphae extending within the FOV as well as all associated host membrane rupture and host cell deaths detected over the course of the experiment can be directly associated with the IYs present within the FOV at time 0. Thus, each FOV constitutes a well-defined ‘infection site’ that can be analyzed in detail at the single-cell level. An additional advantage of this setup is that the low MOI used makes manual and automated detection of discrete infection events technically feasible over many hours without the FOV overcrowding.

### Quantification of host membrane rupture and host cell death during infection of TR146 cells with WT and *ece1*Δ/Δ *C. albicans* strains

We first studied the dynamics of host membrane rupture during the first 12 hours of WT and *ece1*Δ/Δ infection (**Fig 1C, S1 Movie**). Multiple infection sites containing between 1 and 4 IY were imaged every 10 minutes, with individual galectin-3 recruitments recorded at each time point in four independent experiments. Quantification of Gal-3 recruitments was stopped after 10 hours, as past this point infection sites became very crowded resulting in ambiguity in Gal-3 recruitment identification. As expected, during WT infection, total Gal-3 recruitments increased over time, with a lag time of around 200 minutes from the start of infection, while during *ece1*Δ/Δ infection, total Gal-3 recruitments remained low but did show a slight increase over time. *ece1*Δ/Δ hyphal extension and invasion were not attenuated compared to the WT, as previously reported[6]. Infection sites containing a higher number of WT IY exhibited a statistically significant increase in host membrane rupture at 10 hours of infection, a trend not observed for the *ece1*Δ/Δ strain, indicating a correlation between fungal load and host membrane rupture during WT infection (**S2 Fig**). The total number of WT Gal-3 recruitments recorded at 10 hours in sites containing a single IY was on average 12 (n = 3). For *ece1*Δ/Δ the average for sites containing a single IY was 1.3 (n = 3, average for all infection 1.13, n = 57), so that WT infection resulted in around ten times more membrane breaching events compared to *ece1*Δ/Δ infection.

Next, we performed an equivalent analysis on the number of host cell deaths as indicated by individual SYTOX green labeling events during a 24-hour infection with 30 minutes imaging intervals using WT, and *ece1*Δ/Δ strains (IY range 1-4) as well as non-infected control (NI) in three independent experiments (**Fig 1D, S2 movie**). During WT infection, host cell death increased over time, with a lag time of around 10 hours from the start of infection. During *ece1*Δ/Δ infection, total host cell death remained low, but did show a slight increase over time, a trend also observed in the non-infected control. Infection sites containing a higher number of WT IY exhibited an increase in host cell death at 24 hours of infection, though this was not statistically significant (ANOVA p-value: 0.6), a trend not observed for the *ece1*Δ/Δ strain. This result suggests a possible correlation between fungal load and host cell death during WT infection (**S2 Fig**). The total number of WT host cell death recorded at 24 hours in all infection sites was on average 155.3 (IY = 1-4, n = 20). For *ece1*Δ/Δ the average for all infection sites was 25 (IY = 1-4, n = 15) and for the non-infected control 20 (n = 11). As the values of total host cell death at 24 hours during *ece1*Δ/Δ infection are comparable to the non-infected control, *ece1*Δ/Δ infection appears to result in little or no additional host cell death in our system. WT infection results in around six times more host cell deaths compared to non-infected cells or cells infected with *ece1*Δ/Δ.

### Invading hyphae extend in proximity to the microplate surface

Following our global analysis of host membrane rupture and cell death during infection, we turned to examine the architecture of hyphal invasion within our experimental model. We examined the trajectories of invading hyphae using optical z-sectioning with a high-resolution objective (100X) of live and fixed infection sites at around 7 hours post-infection (**Fig 2A, B, C**). Imaging of CM staining of non-fixed samples revealed that invasion pockets formed by invading hyphae are found in close proximity to the microplate surface (i.e. lowest z-section) (**Fig 2A, S3 movie**). Revisiting the exact same site following fixation and staining with Calcofluor white (CFW), a stain used to label *C. albicans* cell wall, and phalloidin-FITC, staining actin, confirmed that hyphae indeed extend along the microplate surface, and appear to ‘cut-through’ the host basolateral cortical actin (**Fig 2B, S4 movie**). In order to reach the microplate surface, hyphae initially invade via a downward-diagonal ‘apical invasion’ path, after which they remain in proximity to the microplate surface for the remainder of infection (**Fig 2C**). While deviations in hyphal trajectories away from the microplate surface were occasionally observed, the overall ‘invasion architecture’ described here was maintained throughout infection in both the WT and *ece1*Δ/Δ strains (**S3 Fig**).

**Fig. 2.**
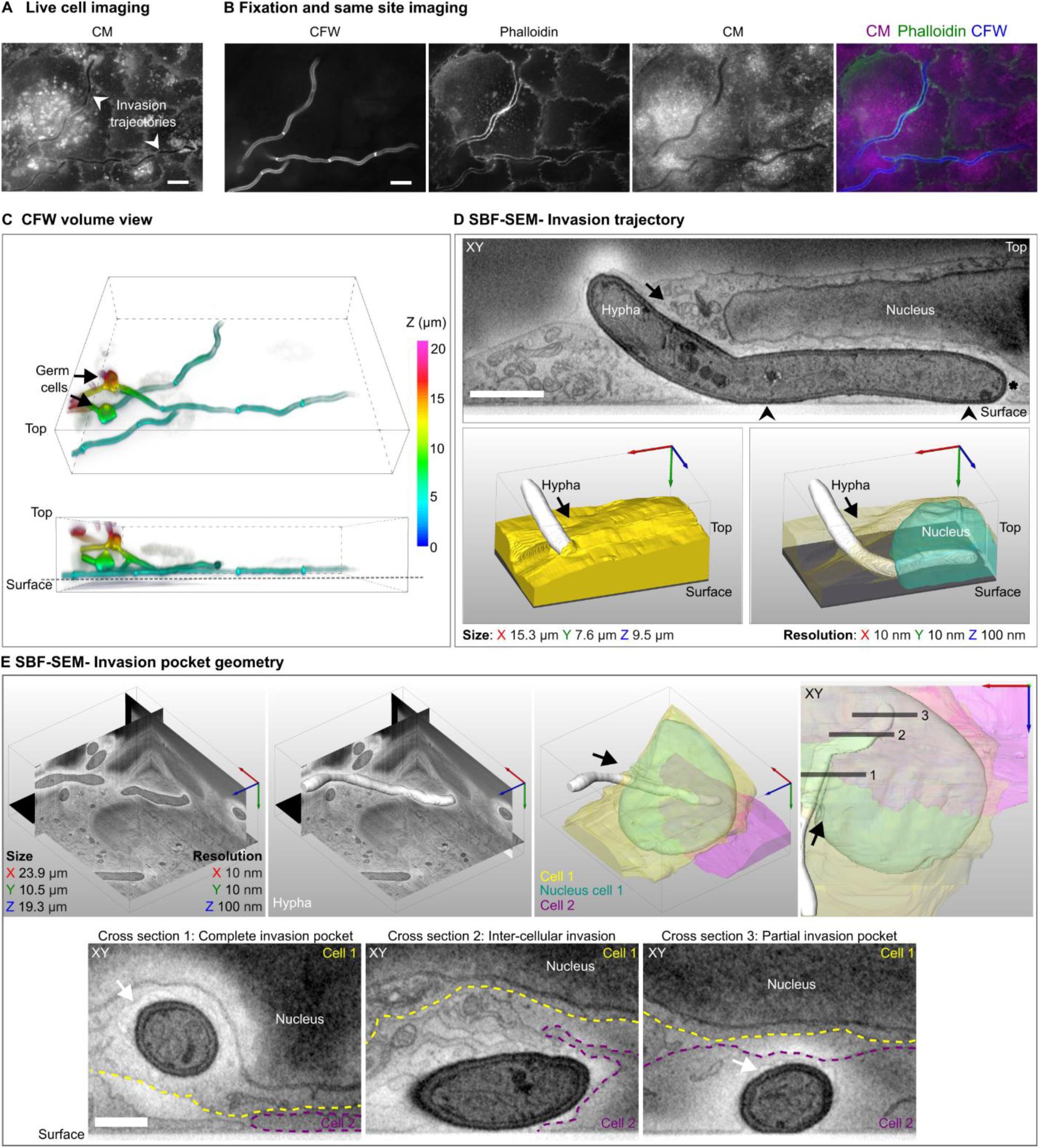
‘Invasion architecture’ of TR146 monolayer infection. (A) CM staining of live (non-fixed) WT invasion site 7 hours post-infection. The optical section (z) nearest to the microplate surface is presented. Two invasions trajectories are observed (white arrowheads). (B) Same-site imaging of fixed samples stained with CFW, a fungal cell wall marker, and phalloidin, an actin marker. CM staining after fixation is presented for reference. In all images the optical section nearest to the microplate surface is presented. (C) A three-dimensional representation of CFW staining in panel B using z-stack color coding. (D) Serial block face-scanning electron microscopy (SBF-SEM) of a WT invasion site 5 hours post-infection. A single XY section is presented (top), with points of direct contact with the microplate surface indicated with black arrowheads and the hyphal apex indicated with an asterisk. Two three-dimensional representations of the same dataset (bottom) showing the epithelial cell surface (yellow) invaded by a hypha (white) (left) and the invasion trajectory within the host cell underneath the host nucleus (teal) (right). (E) SBF-SEM dataset of a hypha (white) invading two host cells (Cell 1-yellow, Cell 2-magenta) in sequence. The nucleus of Cell 1 is also presented (teal). XY cross-sections from within the volume are presented at three different planes along the invasion trajectory: cross section 1-a ‘complete’ invasion pocket surrounding the hypha segment within Cell 1; cross section 2- a hypha segment within the intercellular space between Cell 1 and Cell 2; cross section 3- a ‘partial’ invasion pocket around a hypha segment in direct contact with the microplate surface within Cell 2. The invasion pocket membrane is indicated with a white arrow. In all panels, the direction of hyphal extension is indicated with black arrows. Scale bars are 10 µm in A and B, 2 µm in D and 1µm in E. The dimensions and resolution of the SBF-SEM volumes presented are listed within the figure. See also **S3, S4, S5, S6 Movies.**

### Invasion occurs via intra- and inter-cellular routes and features different invasion pocket geometries

While high-resolution optical z-sectioning analysis revealed the general ‘invasion architecture’ in our system, the resolution limit of light microscopy did not allow us to determine the precise three-dimensional path of invasion within the epithelial layer, epithelial membrane organization around invading hyphae and whether hyphae come into direct contact with the microplate surface during invasion. To address these questions, we employed an advanced volume electron microscopy technique termed ‘serial block face-scanning electron microscopy’ (SBF-SEM), capable of providing three-dimensional nano-scale views of *C. albicans* invasion sites [5]. Two WT invasion sites are presented (**Fig 2D, 2E**), representing typical invasion sites observed in both WT and *ece1*Δ/Δ SBF-SEM volumes acquired five hours post-infection (n = 11 WT, 7 *ece1*Δ/Δ complete invasion sites - i.e. invading hypha entirely captured within the 3D volume and n = 13 WT, 8 *ece1*Δ/Δ incomplete invasion sites - i.e. some invading hypha segments are outside the acquired volume). SBF-SEM analysis revealed that following an ‘apical invasion’ through the first invaded host cell, hyphae can come directly into contact with the microplate surface (**Fig 2D, S5 Movie**). Strikingly, invading hyphae appeared to ‘burrow’ underneath the host nucleus such that it was substantially deformed. Invasion within the nucleus itself or nuclear envelope damage along the hyphal trajectory were never observed. Examination of the nano-scale epithelial layer organization along the path of a longer hypha that invaded two host cells in sequence revealed that invasion can occur both within cells (intra-cellular route) and between cells (inter-cellular route) (**Fig 2E, S6 Movie**). During intra-cellular invasion, we observed both ‘complete’ invasion pockets where host membranes fully encircled the invading hyphae, as well as ‘partial’ invasion pockets, whereby hyphae extending in direct contact with the microplate surface were partially encircled by the plasma membrane of the epithelial cell above. SBF-SEM analysis of *ece1*Δ/Δ invasion sites did not reveal any differences compared to WT invasion sites at this time point (**S3 Fig**). Overall, the ‘invasion architecture’ in our model is characterized by hyphae extending in proximity to microplate surface, where invasion can occur within cells, between cells and in direct contact with the substratum.

### The spatial organization of host membrane rupture sites is regulated by candidalysin and is predictive of host cell death

Next, we examined the spatial organization of host membrane rupture sites, its regulation by candidalysin and correlation with host cell death (**Fig 3**). Analysis of Gal-3 recruitment patterns observed during the first 10 hours of infection in three independent experiments revealed that patterns can be classified into three distinct categories that are not temporally linked (i.e. one pattern does not transition into another over time) (**Fig 3A, S7-9 movies**): ‘tip’ recruitment, defined by Gal-3 recruitment in a small region at the invasion pocket tip (i.e. near the hyphal apex), ‘tubular’ recruitment, defined by Gal-3 recruitment along a short section of the ‘neck’ of the invasion pocket, which excludes the invasion pocket tip, and ‘elongated’ recruitment, defined by a long Gal-3 recruitment along the invasion pocket length that includes both the tip and the neck. Pattern classification was performed manually based on the parameters described above. Quantification of the distribution of recruitment patterns during infection revealed that for the WT strain, ‘tip’ recruitment is observed in 66.8% of all events, ‘tubular’ in 19.6%, and ‘elongated’ in 13.6% (n = 404). For *ece1*Δ/Δ, all events occurred in the ‘tip’ recruitment pattern (n = 23) (**Fig 3B**). Taken together, membrane rupture occurs with three distinct spatial organizations, two of which are dependent on the presence of candidalysin (‘tubular’ and ‘elongated’), and one observed with or without the presence of candidalysin (‘tip’). Past the first 10 hours of infection, infection sites became very crowded, making unambiguous determination of Gal-3 recruitment patterns impractical. Furthermore, from this point on, Gal-3 recruitments resulting from hypha-hypha collisions became increasingly abundant, likely driven by the invasion architecture of our experimental model (see **Discussion).**

**Fig. 3.**
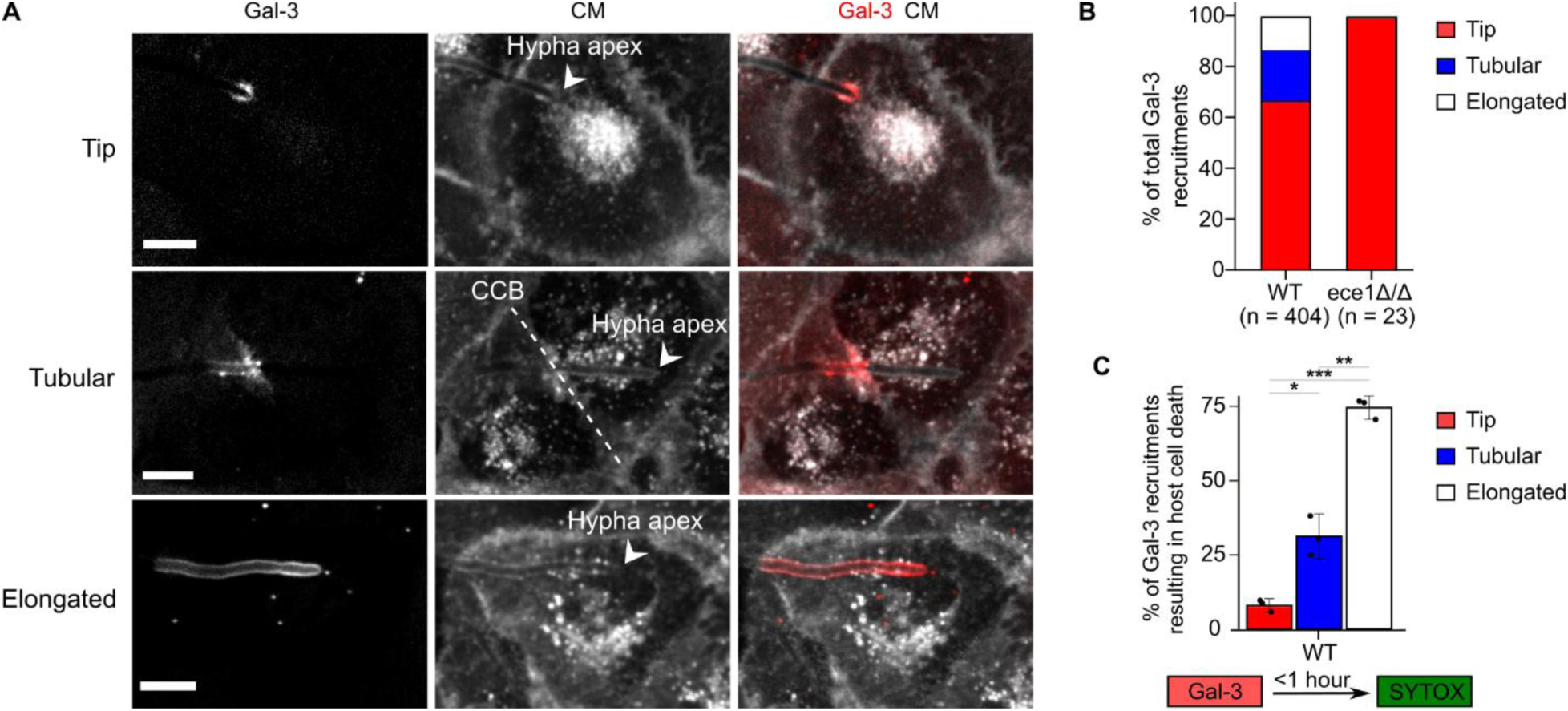
Galectin-3 recruitment patterns are regulated by candidalysin and are predictive of host cell death. (A) Galectin-3 recruitment patterns were classified into three categories: ‘tip’, ‘tubular’ and ‘elongated’. The hypha apex (arrowheads) and cell-cell boundary (CCB, dotted line) are indicated. (B) Distribution of recruitment patterns during WT and *ece1*Δ/Δ infection (n = 404 WT and n = 23 *ece1*Δ/Δ Gal-3 recruitments from three independent experiments, χ2 test for homogeneity p-value **). (C) Correlation between Gal-3 recruitment pattern and host cell death. Gal-3 recruitment was considered correlated to host cell death if a SYTOX signal was observed within the same cell up to one hour after Gal-3 recruitment. As no host cell death was observed during infection with the *ece1*Δ/Δ strain, only WT data is shown (n = 86 SYTOX labeling events following 404 Gal-3 recruitments in three independent experiments, student t-test is presented). Scale bars are 10 µm. p-value thresholds: 0.05 (*), 0.01 (**), 0.001 (***). See also S7-9 movies.

Next, we analyzed the correlation between the three Gal-3 recruitment patterns detected and host cell death as indicated by SYTOX labelling. An individual galectin-3 recruitment pattern was considered correlated to host cell death if a subsequent SYTOX signal was observed within one hour (six acquisition intervals) in the same cell in which Gal-3 recruitment was initially observed. In three independent experiments we found that for the WT strain, ‘tip’ recruitment was correlated to host cell death in 8% of events (n = 21/270), ‘tubular’ recruitment in 30.5% of events (n = 24/79) and ‘elongated’ recruitment in 72% of events (n = 41/55) (**Fig 3C**). Of note, ‘elongated’ recruitment patterns were largely concurrent with SYTOX labelling within the same cell (within the temporal resolution of our system), suggesting they occur in response to a general membrane permeabilization occurring during host cell death, in which Gal-3 is recruited along the entire invasion pocket membrane, a phenomenon previously observed during *C. albicans* HeLa cell invasion[5]. During infection with the *ece1*Δ/Δ strain, we did not observe any host cell death following Gal-3 recruitment (n = 23). These results suggest that candidalysin-dependent Gal-3 recruitment patterns (i.e., observed only during WT infection) are linked to increased cell death compared to the ‘tip’ recruitment pattern, observed in both WT and *ece1*Δ/Δ infections. [5]

### Host membrane rupture occurs at two distinct host subcellular niches: the juxtanuclear region and the cell-cell boundary

Our detailed analysis of Gal-3 recruitment dynamics and spatial organization led to the surprising observation that the location of Gal-3 recruitments within host cells is associated with two specific host subcellular niches: the vesicle rich juxtanuclear region (JNR) and the cell-cell boundary (CCB). This phenomenon is illustrated using a high-temporal and -spatial resolution acquisition (60X objective) of a two-hour WT infection window with imaging intervals of two minutes using the Gal-3 and CM channels (**Fig 4A, 4B, S10 movie**). The invasion path of a single hypha extending through two host cells over the course of around 90 minutes is presented in an overview: at time = 0, the invading hypha is already enveloped in an invasion pocket having previously invaded other host cells; first the hypha extends in proximity to the vesicle rich juxtanuclear region in Cell 1 (JNR1); the hypha then extends past the cell-cell boundary (CCB) separating the two host cells; and finally the hypha extends near the juxtanuclear region of Cell 2 (JNR2) (**Fig 4A**). Gal-3 recruitments occur several times along this path of invasion, with the first event occurring in proximity to JNR1 (Rupture site 1-R1), followed by a rupture at the CCB (Rupture site 2 - R2), and finally in proximity to JNR2 (Rupture site 3 - R3) (**Fig 4B**). Insets show the spatial organization of each recruitment, with ‘tip’ recruitments for R1 and R3 and ‘tubular’ recruitment for R2.

**Fig. 4.**
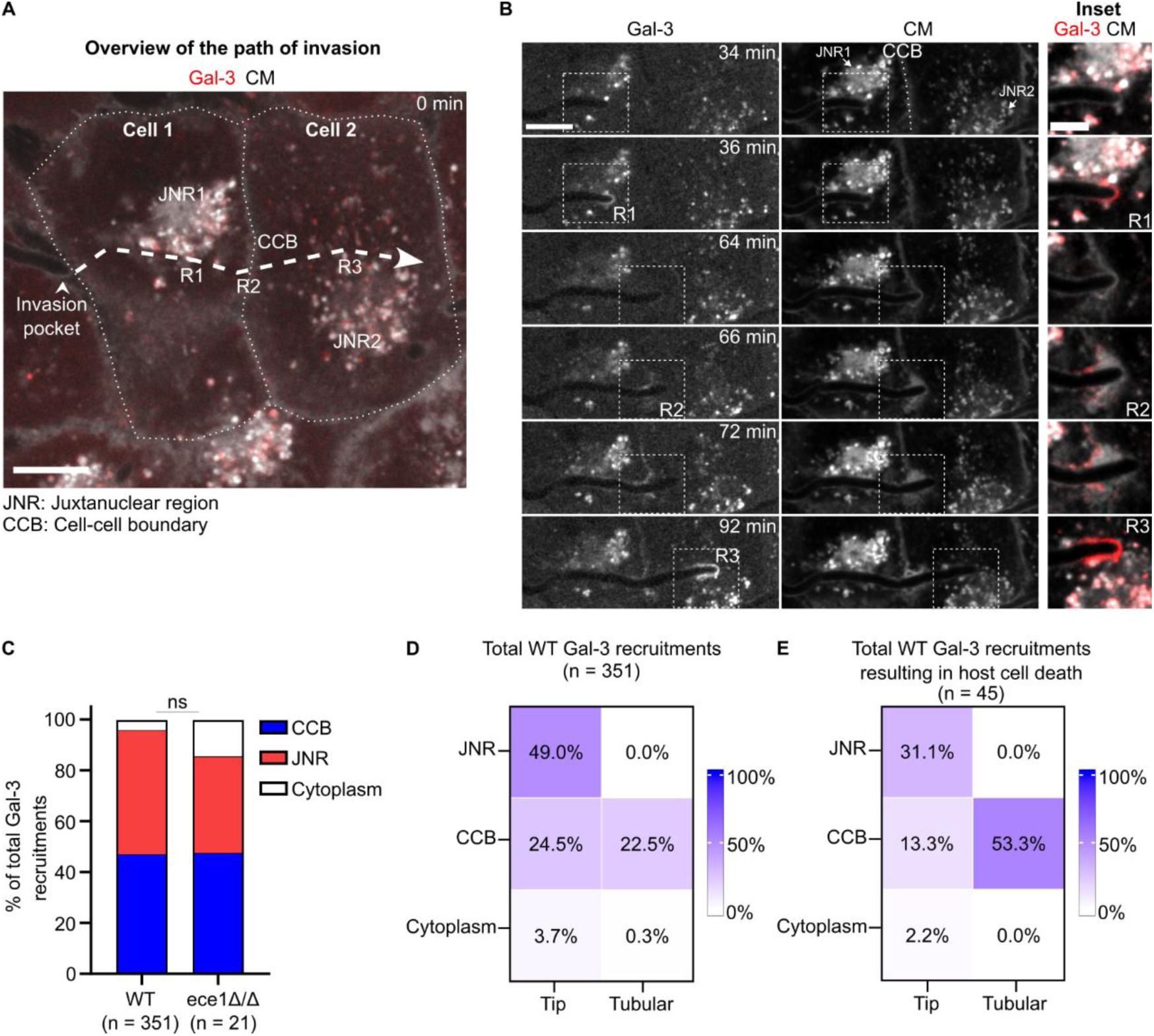
Gal-3 recruitments are localized to two host subcellular niches. High temporal- and spatial-resolution live cell imaging showing multiple galectin-3 recruitments occurring along the invasion path of a single hypha invading two host cells in sequence. (A) An overview of the hypha trajectory (dashed line) starting at 0 min is presented. Cell 1 and Cell 2 are outlined with a dotted line. An invasion pocket labelled by CM is already present at t = 0 (arrowhead). The hypha trajectory traverses the juxtanuclear region of Cell 1 (JNR1), the cell-cell boundary (CCB) and the juxtanuclear region of Cell 2 (JNR2) with three membrane rupture events (R1, R2, R3). (B) A detailed view of Gal-3 recruitments occurring at JNR1 (R1), the CCB (R2), and JNR2 (R3) is presented, with insets showing a magnified view. Imaging was performed for 2 hours with 2 minutes intervals. (C) Quantification of Gal-3 recruitment location, classified into three categories: cell-cell boundary (CCB), juxtanuclear region (JNR) and cytoplasm. Analysis is based on 12-hour infection acquisitions with an imaging interval of 10 min from three independent experiments (n = 351 WT and n = 21 *ece1*Δ/Δ Gal-3 recruitments). The difference in galectin-3 recruitment locations between WT and *ece1*Δ/Δ infections is not statistically significant (χ2 test). (D) Distribution of WT Gal-3 recruitment ‘pattern-location’ pairs (n = 351 Gal-3 recruitments in three independent experiments. χ2 test p-value ***) (E) Distribution of WT Gal-3 recruitment ‘pattern-location’ pairs resulting in host cell death (n = 45 Gal-3 recruitments resulting in host cell death, indicated by SYTOX labeling within 1 hour of Gal-3 recruitment, χ2 test p-value ***). Scale bars are 10 µm and 5 µm in inset. p-value thresholds: 0.05 (*), 0.01 (**), 0.001 (***). See also S10 movie.

Quantitative analysis was performed via manual annotation of the location of each Gal-3 recruitment observed during three independent 12-hour infection acquisitions using both WT and *ece1*Δ/Δ strains (n = 351 WT and n = 21 *ece1*Δ/Δ). Locations were classified into three categories: recruitment at the CCB, recruitment in proximity to the JNR, and recruitment at the cytoplasm (i.e., without proximity to the JNR or CCB) (**Fig 4C**). As ‘elongated’ Gal-3 recruitment patterns (accounting for 13.6% of all recruitments) span the length of most of the host cell, their location could not be classified unambiguously, and thus they were omitted from the location analysis. For the WT strain, 47% (165/351) of Gal-3 recruitments occurred at the CCB, 49% at the JNR (180/351), and 4% (14/351) at the cytoplasm. For the *ece1*Δ/Δ strain 48% (10/21) of Gal-3 recruitments occurred at the CCB, 38% (8/21) at the JNR, and 14% at the cytoplasm (3/21). The distribution of locations between the two strains was not statistically different. As a reference, we estimate that the cytoplasm (including the thin cytoplasmic area between the basolateral membrane and the nucleus above) occupies around 87% of the cell area in proximity to the microplate surface, compared to 13% occupied by the JNR (**S4 Fig**). Taken together, our results suggest that host membrane rupture occurs predominantly at two distinct host subcellular niches: the CCB and the JNR, and that membrane rupture location is not regulated by candidalysin.

### Membrane rupture at the cell-cell boundary is the main driver of host cell death

Next, we combined the Gal-3 recruitment pattern analysis (**Fig 3B**) with the Gal-3 recruitment location analysis (**Fig 4C**), assigning both a recruitment pattern and a location to WT recruitments (n = 351 Gal-3 recruitments, ‘elongated’ recruitment pattern excluded, see above) as ‘pattern-location’ pairs (**Fig 4D**). *ece1*Δ/Δ infection was not analyzed as all recruitments occurred with the ‘tip’ pattern. Our results revealed several interesting trends: almost half of all recruitments (49%) occurred as ‘tip-JNR’, whereas ‘tubular-JNR’ pairs were never observed. In other words, only ‘tip’ recruitments were observed at the JNR region. Tubular recruitments occurred almost exclusively at the CCB (79/80 total tubular Gal-3 recruitments), suggesting this recruitment pattern is strongly associated with hyphae traversing the cell-cell boundary. ‘Tip’ and ‘tubular’ recruitments at the CCB occurred in approximately equal proportions (24.5% and 22.5% of all recruitments respectively).

Finally, WT ‘pattern-location’ pairs were correlated with subsequent host cell death (**Fig 4E**), using the criteria of SYTOX signal appearing within one hour of Gal-3 recruitment in the same cell (see also **Fig 3C**). Overall, 13% of total Gal-3 recruitments (‘elongated’ pattern excluded) resulted in host cell death within one hour (n = 45/351). Within this subset (n = 45), 53.3% of host cell deaths occurred following ‘tubular-CCB’ Gal-3 recruitment. As ‘tubular-CCB’ recruitments account for only 22.5% of all Gal-3 recruitment (**Fig 4D**), we conclude that the ‘tubular-CCB’ pair is strongly correlated with host cell death. In contrast, 31.1% of host cell deaths occurred following ‘tip-JNR’ Gal-3 recruitment, while accounting for 49% of total Gal-3 recruitments. Thus, we conclude that the ‘tip-JNR’ pair is weakly correlated to host cell death. Comparing the correlation between Gal-3 recruitment location and host cell death (irrespective of the recruitment pattern) reveals that while the proportion of JNR and CCB Gal-3 recruitments is nearly identical (49% to 47% respectively, **Fig 4D**), CCB recruitments are correlated to 66.6% of total host cell death, versus only 31.1% of recruitments in the JNR. Taken together, Gal-3 recruitments at the CCB, and in particular those associated with the ‘tubular’ recruitment pattern, are strongly correlated to subsequent host cell death, suggesting membrane rupture at the CCB is the main driver of subsequent infection damage. In contrast, membrane rupture at the JNR (associated exclusively with ‘tip’ recruitment pattern) is weakly correlated to host cell death, suggesting this subcellular niche is relatively non-damaging, and plays a smaller role in generating long-term infection damage.

### High temporal- and -spatial resolution imaging of membrane rupture reveals niche-specific attributes

To obtain more detail about the process of membrane rupture in each subcellular niche, we employed high temporal- and spatial-resolution live cell imaging (60X objective) over a two-hour infection window when Gal-3 recruitments were abundant, with two minutes acquisition intervals (**Fig 5**). We did not observe any transitions over time between the different recruitment patterns (tip, tubular, elongated) even at high temporal-resolution, confirming these patterns are not temporally linked and thus represent distinct outcomes. A typical ‘tubular/CCB’ Gal-3 recruitment is presented in detail (**Fig 5A, S11 movie**): at the start of the timeline (0 min), a hypha enveloped by an invasion pocket has already extended past the length of the first cell (Cell 1) with the hypha apex at the CCB. In the next minutes (8, 16 min) the hypha extends into the second cell (Cell 2). Next, as the hypha continues to extend, a tubular Gal-3 recruitment appears (24 min). Notably, Gal-3 recruitment in the ‘tubular’ pattern is observed only within the first invaded cell. Gal-3 labelling is also observed in a small section of the membrane orthogonal to the invasion pocket, suggesting damage is not limited only to the invasion pocket membrane itself. Finally, hyphal extension within an invasion pocket continues in Cell 2. This ‘cell selective’ CCB recruitment pattern was commonly observed (see also **Fig 3A** and **Fig 4B**, for additional examples). Overall, our data suggests that upon contact with the CCB, hyphae ‘push’ the invasion pocket membrane formed in the first invaded cell into the second invaded cell, until a ‘critical point’ is reached when membrane rupture occurs selectively in the first invaded cell only (see **Discussion**).

**Fig. 5.**
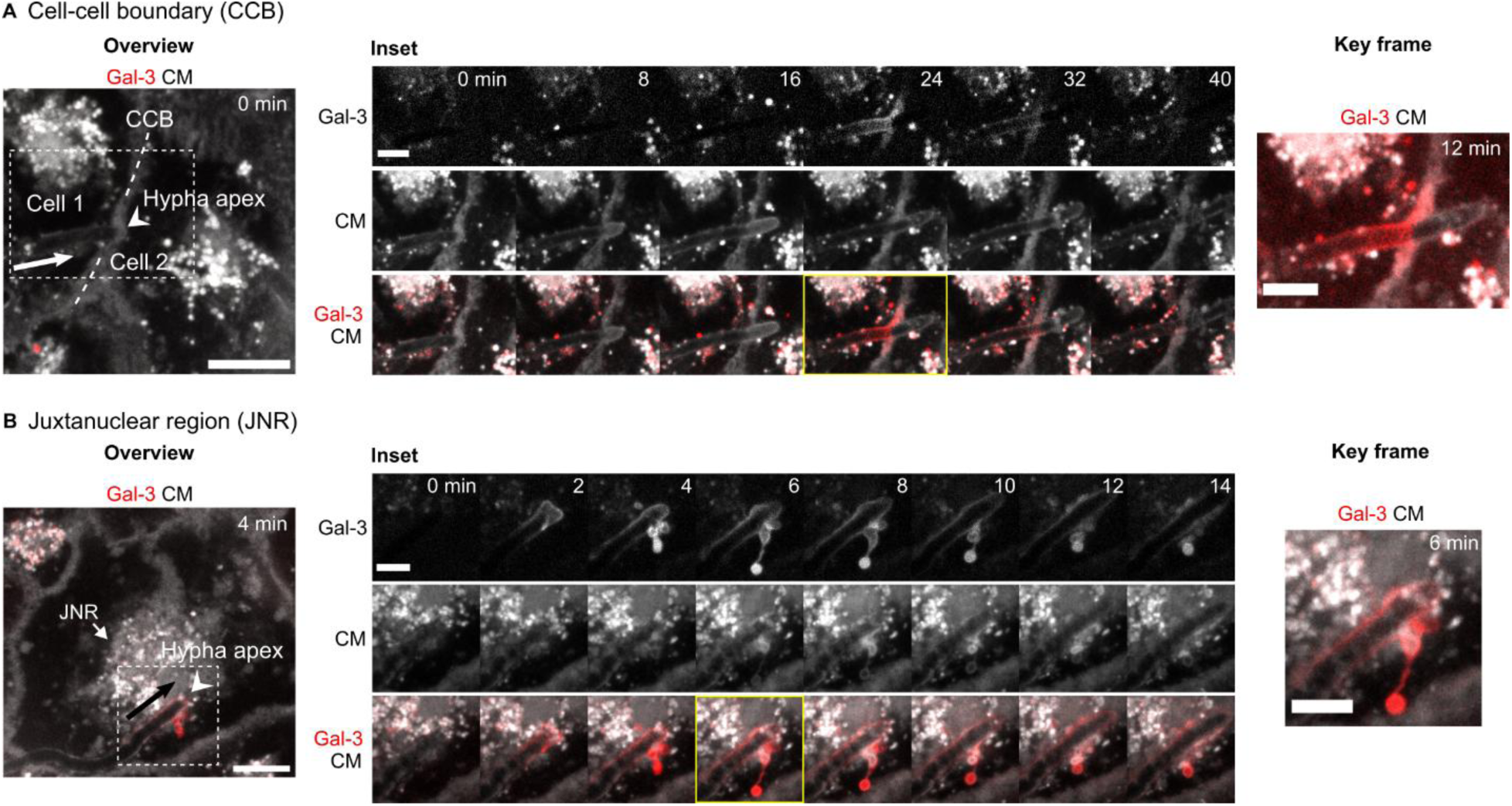
Specific attributes of host membrane rupture at the CCB and JNR. High temporal- and spatial-resolution live cell imaging showing rupture dynamics and organization specific to each subcellular niche (A) ‘tubular/CCB’ Gal-3 recruitment. An overview shows the invasion pocket membrane at the CCB between Cell 1 and Cell 2 at t = 0 min. The hypha apex is indicated (arrowhead). The direction of extension is noted with an arrow. Inset (from dotted frame) shows the invasion pocket extending into Cell 2 followed by a tubular Gal-3 recruitment limited to Cell 1. The invasion pocket then continues to extend into Cell 2. A key frame (yellow) highlights ‘cell selective’ Gal-3 recruitment. See also **S11 movie.** (B) ‘tip/JNR’ Gal-3 recruitment. An overview shows an invasion pocket traversing the JNR. The hypha apex is indicated (arrowhead). The direction of extension is noted with an arrow. Inset (from dotted frame) shows Gal-3 tip recruitment associated with vesicle-like structures. A key frame (yellow) highlights the Gal-3 positive vesicles and tether. See also **S12 movie**. Scale bars are 10 µm in overviews and 5 µm in insets and key frames.

Next, we examined Gal-3 recruitment patterns associated with JNR rupture in detail. A typical ‘tip/JNR’ Gal-3 recruitment is presented (**Fig 5B, S12 movie**). The first time point in which Gal-3 recruitment is observed shows a typical ‘tip’ recruitment pattern. Closer examination of this structure suggests the presence of two vesicle-like structures at the hypha apex, that are seen connected to the rest of the invasion pocket membrane (2 min). Next, several vesicles labelled with Gal-3 ‘pinch off’ the enveloping membrane but remain close to it (4 min). A single vesicle moves away from the invasion pocket membrane, but remains tethered to it via a thin membrane connection (6 min) at which point it is ‘pulled back’ (8, 10, 12 min). As the hypha and invasion pocket continue to extend, Gal-3 positive vesicles are ‘left behind’ and remain in a static location (12, 14 min). The formation and ‘removal’ of Gal-3 positive vesicles were commonly observed at JNR membrane rupture sites, suggesting they represent a ‘niche-specific’ attribute. While in this study we did not attempt to further characterize the precise nature of these vesicles, our observations are in good agreement with a previous study of *C. albicans* TR146 cell infection that demonstrated the presence of membrane-derived blebs at sites of membrane rupture, suggested to be part of a specific host membrane repair response to *C. albicans* invasion (see **Discussion**)[14].

## Discussion

In this study, we employed live cell imaging and volume electron microscopy to study the interplay between hyphal extension, candidalysin secretion, and host damage during *in vitro* epithelial invasion by *C. albicans*. We quantify host membrane rupture and cell death over the course of infection at the single cell level for both WT and *ece1*Δ/Δ strains. We also determine the ‘invasion architecture’ in our experimental model, showing that following an initial ‘apical’ invasion, hyphae extend in proximity to the microplate surface and can invade the epithelial layer via intra- and inter-cellular routes. We show that membrane rupture is spatially organized in three distinct patterns that are differentially regulated by candidalysin. Strikingly, we observe that membrane rupture occurs at two host subcellular niches: the cell-cell boundary (CCB) and the juxtanuclear region (JNR), and that this distribution is candidalysin-independent. Our analysis also reveals that rupture spatial organization, location and subsequent host cell death are linked, suggesting CCB rupture is strongly correlated to subsequent host cell death. Finally, we reveal two ‘niche-specific’ attributes-cell selective rupture at the CCB and vesicle formation and removal at the JNR.

Previous studies have compared host damage inflicted by WT and *ece1*Δ/Δ strains in different models, providing an indication of the relative ‘contribution’ of candidalysin to total infection damage[11,20]. LDH release assays of TR146 cells after 24 hours of infection showed approximately 8-fold decrease in damage when using the *ece1*Δ/Δ strain compared to the WT strain[6]. This result is in general agreement with our study, which indicates a 10-fold decrease in host membrane rupture at 10 hours of infection and a six-fold decrease in host cell death at 24 hours of infection using the *ece1*Δ/Δ strain compared to the WT strain. Thus, our study further supports the well-established role of candidalysin as a key fungal factor regulating host damage during infection.

We show that *C. albicans* invasion and damage in our epithelial monolayer model occur via a distinct ‘invasion architecture’ that has important implications for the interpretation of our results. Following an initial apical invasion through the epithelial layer, hyphae extend near the microplate surface for the remainder of infection, revealing a clear limitation of our monolayer model compared to the natural setting where *C. albicans* invade a three-dimensional stratified oral epithelium. Within this setting, invading hyphae were observed extending between and within host cells, and also came into direct contact with the microplate surface itself. Hyphae invading within host cells were thus either completely or partially enveloped by host membranes along their trajectory, depending if invasion occurred within the epithelial layer proper or in direct contact with the surface. While ‘complete’ invasion pockets appear to represent well ‘true’ intra-cellular invasion, ‘partial’ invasion pocket may not be considered truly ‘intra-cellular’, but rather as a specific form of invasion that impacts epithelial membrane organization more locally. Overall, invading hyphae maintained a general ‘downward’ (i.e. extension towards the microplate surface) trajectory, in accordance with the previously described hyphal thigmotropic response to the local environment [21,22]. Indeed, the reported asymmetrical ‘nose-down’ tip morphology of *C. albicans* hyphae, by which hypha polarity machinery and associated growth site are positioned close to the substratum, allowing hyphae to follow contours and probe underlying unconformities in the growth surface matches well with the invasion architecture observed in this study[23]. This architecture stands in contrast to our previous study of invasion into Caco-2 monolayers, where following apical invasion, hyphae extended within the epithelial layer without reaching the microplate surface for hours, traversing multiple host cells within trans-cellular tunnels[5]. The underlying mechanism driving different ‘invasion architectures’ in different epithelial cell types remains unknown, though it may be linked to variations in the layer thickness and internal cellular organization as well as cell-type specific differences in cellular junctions and epithelial membranes. Indeed, infection ‘damage profile’ also shows a clear link with epithelial cell-type, with early damage observed in TR146 infection compared to tunneling and late damage in Caco-2 infection [4,5]. Finally, the invasion architecture in our experimental model likely underlies two other observations made in this study: first, we did not observe by live cell imaging any interactions between invading hyphae and host nuclei such as hyphal change of direction or membrane rupture, despite nuclei representing the most rigid part of the epithelial cell. Volume electron microscopy revealed that invading hyphae can drastically deform the shape of host nuclei while maintaining hyphal trajectory. Additionally, hyphae were never observed within nuclei, and we did not detect any nuclear envelope damage associated with invading hyphae. This suggests that the nuclear envelope does not rupture but instead deforms in response to hyphal extension. This phenomenon was previously observed during *C. albicans* invasion of HeLa and Caco-2 cells, suggesting it is not cell-type dependent[5]. If and how nuclear deformations impact host cells during invasion remains to be studied. Second, we observed that around ten hours post-infection, hypha-hypha collisions become a dominant cause of host membrane rupture. The invasion architecture of our system would imply that at later time points, all invading hyphae are extending near the microplate surface in a near ‘two-dimensional space’, therefore increasing the chances of hyphal collisions as the number and length of hyphae increases over time. As hyphal collisions are expected to be a much rarer event when hyphae extend within a three-dimensional stratified epithelium, the physiological relevance of infection damage measured at late time points in our model is brought into question. Future studies in three-dimensional multi-layered infection models such as reconstituted epithelia[24], should allow to directly address the importance of model dimensionality for studying *C. albicans* infection *in vitro*.

Several studies have demonstrated that damage levels caused by the direct addition of exogenous candidalysin peptide to epithelial cells is significantly lower than damage observed during *C. albicans* infection[6,11]. Furthermore, infection with a yeast-locked *C. albicans* strain engineered to secrete candidalysin also results in low levels of damage compared to filamenting cells[7]. These results point towards a requirement for hyphal invasion and candidalysin secretion to act together to inflict maximal damage during infection. To explain this requirement, it has been proposed that the invagination of the host plasma membrane into an invasion pocket by invading hyphae creates a microenvironment that facilitates candidalysin accumulation over time until a lytic concentration of the toxin is reached, followed by damage to the invasion pocket membrane[6,25]. Thus, the invasion pocket enables physiological concentrations of candidalysin to locally reach the level required for membrane lysis, matching the lytic concentrations observed when candidalysin is added exogenously. While candidalysin has been shown to be present along the invasion pocket length, the dynamics and mechanism driving candidalysin accumulation remain unknown[7]. As the proposed ‘accumulation’ mechanism suggests that candidalysin concentration within the invasion pockets reaches a ‘critical’ lytic threshold, one could predict that host membrane rupture during invasion would occur randomly along the path of invasion when this threshold is reached, or alternatively, that longer hyphae should exhibit more membrane damage compared to shorter hyphae, as longer hyphae allow more time for candidalysin accumulation within the invasion pocket [14]. Our study revealed that membrane rupture occurs almost exclusively when hyphae traverse the JNR and the CCB. While we did not directly measure the hyphal length at the time of rupture, we rarely observe (4% of total WT rupture events) membrane breaching occurring as hyphae traverse the host cytosol, despite this being by far the compartment with the largest area near to the microplate surface where hyphae extend, which we estimate at around 87% in TR146 cells. If membrane rupture occurred randomly along the path of invasion, most membrane rupture would be expected to occur during traversal of the cytosol. Additionally, if membrane rupture was dependent on hyphal length, most hyphae would statistically reach the required length for the ‘critical’ lytic threshold while traversing the host cytosol. Both these scenarios stand in stark contrast to our observations. Instead, our results point towards an alternative mechanism linking hyphal invasion and candidalysin activity together, whereby in addition to candidalysin secretion, a second factor, hyphal traversal of distinct subcellular niches, is required to induce membrane rupture and long-term infection damage.

We thus propose the following ‘sequential’ damage mechanism to explain *C. albicans* epithelial damage in our experimental system: Infection begins with hyphal extension along the epithelial layer surface, followed by an apical invasion in which hyphae traverse the epithelial layer (**Fig 2**). From this point onwards, an ‘invasion cycle’ is repeated as each hypha extends in proximity to the microplate surface (**Fig 6**). Candidalysin secretion by an invading hypha surrounded by an invasion pocket membrane (‘complete’ or ‘partial’) results in the formation of a weakened ‘sub-lytic’ invasion pocket along the path of invasion (**Fig 6A**). Next, the hypha may traverse in proximity to the JNR, where the ‘sub-lytic’ invasion pocket is detected by the host, followed by membrane rupture and vesicle removal, possibly as part of a host membrane repair response (**Fig 6B**). Eventually, the extending hypha arrives at the CCB, which acts as a rigid barrier that can exert a counter-force against hyphal extension and the enveloping invasion pocket membrane (**Fig 6C**). CCB traversal leads to rupture of the ‘sub-lytic’ invasion pocket membrane, which in turn is correlated to host cell death and long-term infection damage. Finally, the hypha traverses the CCB towards a neighboring cell forming a new invasion pocket, thus restarting the invasion cycle.

**Fig. 6.**
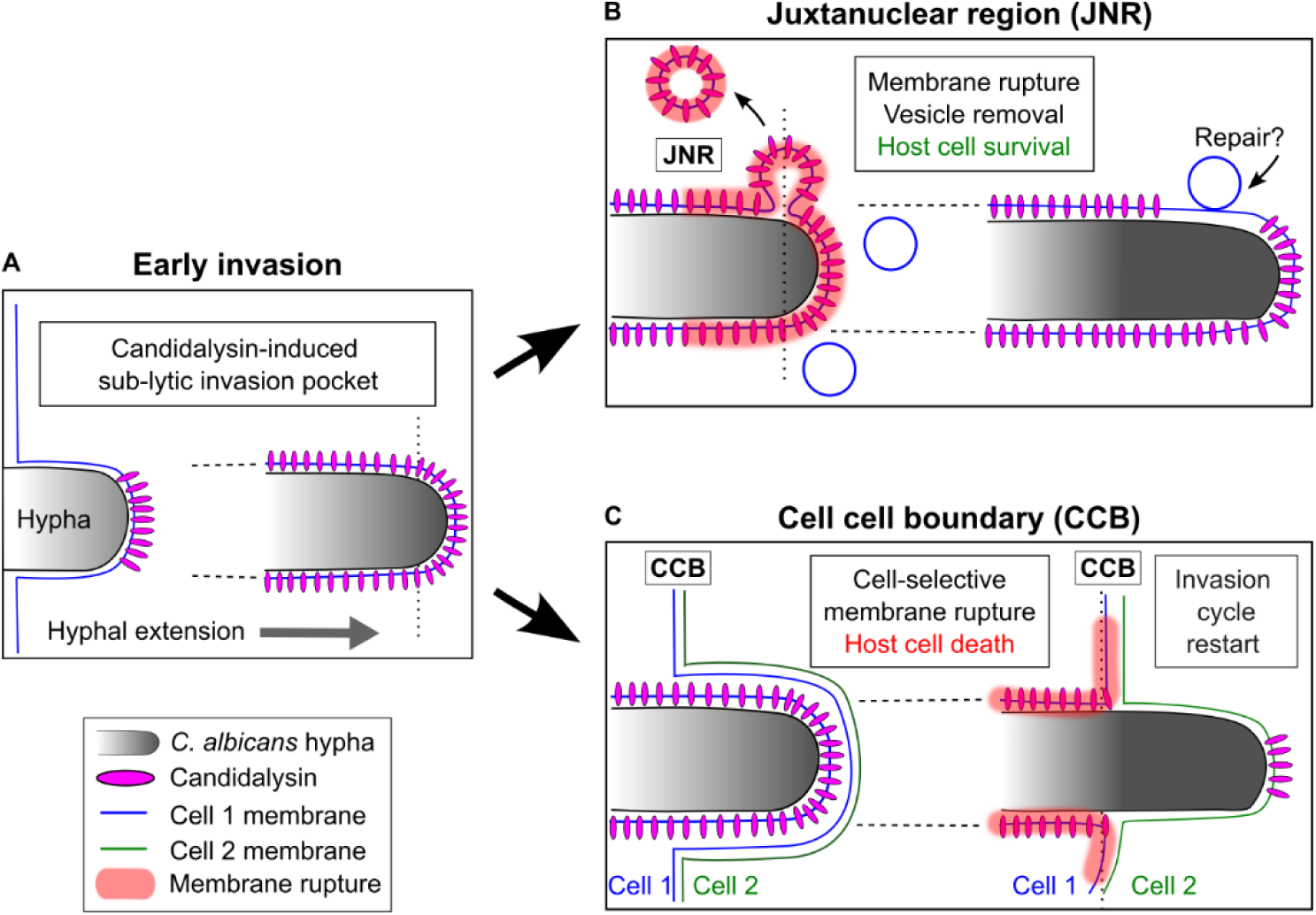
‘Sequential’ damage mechanism for *C. albicans* epithelial infection. An extending hypha initially invades the epithelial layer via an ‘apical invasion’, followed by extension in proximity to the microplate surface where host cells are invaded in sequence (see Fig 2). (A) During early invasion, candidalysin secretion by the invading hypha results in the formation of a ‘sub-lytic’ invasion pocket along the path of invasion prior to membrane rupture. (B) Hyphal traversal of the juxtanuclear region (JNR) results in host membrane rupture followed by removal of vesicles containing ruptured membranes, possibly as part of a regulated host membrane repair response. (C) Hyphal traversal of the cell-cell boundary (CCB) results in cell-selective membrane rupture of the first invaded host cell (Cell 1). The hypha then extends into the second invaded cell (Cell 2), restarting the invasion cycle. Membrane rupture at the CCB is strongly correlated with subsequent host cell death compared to membrane rupture at the JNR, suggesting that CCB traversal is the main driver of long-term infection damage.

In the ‘sequential’ damage mechanism, prior to membrane rupture, invading hyphae are enveloped by a candidalysin-induced ‘sub-lytic’ invasion pocket, with the transition from ‘sub-lytic’ to ‘lytic’ (i.e. membrane rupture), only occurring during hyphal traversal of the JNR and the CCB. In the absence of candidalysin secretion (i.e. during infection with the *ece1*Δ/Δ strain), the number of membrane rupture events is dramatically reduced, though they still occur principally at the JNR and the CCB. This indicates that these niches are intrinsically linked to host membrane rupture, while candidalysin is required to significantly upregulate (10-fold increase) rupture frequency at these sites. Additionally, ‘destructive’ (i.e. highly correlated to subsequent host cell death) ‘tubular’ and ‘elongated’ recruitment patterns are only observed in the presence of candidalysin, suggesting it specifically increases membrane rupture damage-potential. Thus, candidalysin upregulates both the frequency and damage-potential of host membrane rupture, but does not regulate rupture localization. We thus propose that candidalysin secretion and integration into host membranes acts to uniformly weaken the invasion pocket membrane along the path of invasion, resulting in increased rupture frequency and damage when hyphae traverse the JNR and the CCB. Interestingly, nanobody labelling of candidalysin within the invasion pocket of TR146 cells showed it is found in puncta spread along the entire path of invasion, suggesting its integration into host membranes may be uniformly distributed [7]. Conceptually, while the candidalysin ‘accumulation’ mechanism proposed by Moyes et al.[6] represents a ‘one-component’ mechanism dependent only on candidalysin concentration for inducing damage, the ‘sequential’ damage mechanism proposed here represents a ‘two-component’ mechanism, requiring both the action of candidalysin, and hyphae extending through distinct host subcellular niches to induce damage. In the latter mechanism, the function of candidalysin as a toxin may be thought of as ‘priming’ host membranes for subsequent rupture at the CCB and JNR.

Different hypotheses could explain how candidalysin acts to weaken the invasion pocket membrane prior to niche-specific rupture: it is possible that candidalysin integration into the invasion pocket membrane causes only a small scale/local disruption, which is subsequently enhanced during JNR or CCB traversal, leading to large scale disruption and rupture. As candidalysin is a pore forming toxin, it is possible that initial pore damage is limited and lies below the threshold of Gal-3 detection. These pores may then ‘stretch’, enlarge or fuse together during niche traversal, possibly due to increased membrane tension, resulting in membrane rupture[6]. Alternatively, the ‘sub-lytic’ invasion pocket membrane may include precursor states of candidalysin assembly prior to complete pore formation, such as those reported by Russell et al.[10], which in turn mature into fully functional ‘lytic’ pores during JNR or CCB traversal, leading to membrane rupture. Further research is required to address these and other hypotheses underlying candidalysin activity and host damage during *C. albicans* epithelial infection.

As alluded to above, the ‘sequential’ damage mechanism provides an alternative explanation for the requirement of both hyphal invasion and candidalysin secretion to induce full damage potential during infection: hyphal extension is required to drive ‘sub-lytic’ invasion pockets towards the JNR or the CCB, where membrane rupture occurs, a process that cannot be fully replicated by the exogenous addition of candidalysin peptide or by secretion of candidalysin by yeast-locked strains. The ‘sequential’ damage mechanism may also provide an alternative explanation for the dual innate immune response observed during *C. albicans* epithelial infection, though this remains to be experimentally established. This response, proposed to be driven by a concentration-dependent dual-functionality of candidalysin[12], may be attributed instead to the ‘sub-lytic’ invasion pocket driving initial pathogen recognition before appreciable damage, followed by traversal of the JNR and CCB resulting in membrane rupture driving a membrane damage response.

Our work reveals that the CCB is the main site for membrane rupture correlated to subsequent host cell death, suggesting that hyphal traversal of the CCB is the main driver of long-term infection damage. This result is in line with a previous study of *C. albicans* invasion into non-confluent HeLa cells, which demonstrated that a large portion of membrane rupture events (58%) occurred during hyphal traversal of the cell boundary, primarily resulting in host cell death[5]. In contrast, the same study showed that CCB traversal during Caco-2 invasion does not lead to membrane rupture or host cell death at early stages of infection, indicating that different epithelial cell types respond differently to CCB traversal, for reasons that are currently not known. The CCB includes various junctional complexes (adherens junctions, desmosomes, gap junctions and tight junctions) that anchor transmembrane proteins to the cell cytoskeleton (via actin and intermediate filaments) and to neighboring cells[26]. Thus, the CCB represents a mechanically ‘stiff’ barrier that appears to counter the extension of hyphae enveloped by an invasion pocket, resulting in membrane rupture. CCB traversal by the *ece1*Δ/Δ strain rarely results in membrane rupture and never in host cell death, implying that the ‘counter force’ exerted by the CCB, which likely leads to increased membrane tension, is sufficient to rupture sub-lytic invasion pocket membranes, but not native (without candidalysin) membranes. We have also observed that CCB rupture is ‘cell-selective’, damaging only the first invaded cell of the two cells found on each side of the CCB. While the invasion pocket of the first invaded cell at the time of CCB traversal is in a sub-lytic state (having been exposed to candidalysin), membranes of the second invaded cell do not come into direct contact with candidalysin until the invasion pocket of the first cell is ruptured, and are thus ‘native’ during CCB traversal, likely explaining the cell-selective nature of rupture in this niche. Hyphal extension itself is not blocked by the CCB, a result in line with previous studies demonstrating hyphae can invade substrates with stiffness of between 100-200 kPa compared to the stiffness of epithelia, estimated at 40-70 kPa[27]. Of the different junctional complexes, adherens junctions are major determinants of tissue mechanics[28] and were shown to play a key role in *C. albicans* epithelial infection. The *C. albicans* invasins Als3 and Ssa1 have been shown to bind e-cadherin to facilitate induced endocytosis[29,30], while *C. albicans*-secreted aspartic proteinases such as Sap5 can degrade e-cadherin[31]. Indeed, cleavage of c-cadherin has been proposed as one of the mechanisms driving disruption of the intestinal epithelial barrier, resulting in both intracellular and extracellular cleavage events[32]. In addition to adherens junctions, tight junctions also play a role in regulating *C. albicans* invasion of the intestinal barrier[33,34]. Finally, in Caco-2 cells, NFκB activation during infection was shown to stabilize cell-cell junction proteins like E-cadherin, limiting paracellular translocation, while NFκB inhibition allowed hyphae to break down the intercellular barrier in a candidalysin-independent manner[35]. If and how CCB membrane rupture is linked to the various *C. albicans*-cell junction interactions described previously remains to be explored.

Studies in TR146 cells and HeLa cells have shown the presence of a regulated host repair response following *C. albicans* induced membrane damage. This response takes multiple forms including Ca^2+^-dependent autophagic machinery recruitment to the site of damage, ESCRT-dependent membrane blebbing and lysosomal exocytosis[14,36]. More specifically, Westman et al. have shown that during infection of TR146 cells, staining with the lipophilic dye FM4-64 reveals the presence of membrane blebs near the hypha apex which are formed in a candidalysin-dependent manner. As the signal within the blebs was higher than that of the invasion pocket membrane, it was proposed that the observed blebs have undergone membrane rupture. Labelling with nanobodies against candidalysin further revealed the presence of candidalysin within the blebs. The authors suggest that Ca^2+^ influx triggered by candidalysin leads to bleb formation, which in turn allows shedding of the toxin along with the blebbing to terminate the Ca^2+^ influx. Additionally, blebs were shown to be associated with the recruitment of ALG-2, ALIX and ESCRT-III, suggested to constrict and extrude the damaged membrane, allowing eventual membrane repair alongside blebbing. Finally, the authors demonstrate the presence of lysosomes at the hyphal apex, and propose that lysosome exocytosis can serve a complementary role in mitigating candidalysin-induced membrane damage, though lysosome recruitment was also observed in cells infected with the *ece1*Δ/Δ mutant. In this study we observe that JNR rupture is often associated with the formation of Gal-3 labelled vesicles, which are subsequently removed from the site of damage. These observations are consistent with the study by Westman et al., and suggest that JNR rupture and associated vesicle dynamics are part of a regulated host membrane repair response. Indeed, the JNR of TR146 cells was shown to contain lysosomes, further supporting this suggestion[37]. It is possible that host factors located in JNR vesicles detect the sub-lytic invasion pocket as hyphae traverse the JNR, followed by membrane rupture. The precise nature of the signaling involved in JNR rupture is not known, though one possibility is that ‘sub-lytic’ invasion pockets generate a Ca^2+^ influx[14], which in turn is detected by Ca^2+^-dependent receptors found in vesicles or the endoplasmic reticulum in the JNR[38]. Overall, the specific attributes associated with JNR rupture (vesicle formation and removal), the weak correlation of JNR rupture with host cell death compared to CCB rupture, and the ‘non-destructive’ ‘tip’ Gal-3 recruitment pattern associated with this niche, all point towards JNR rupture acting as part of a regulated host membrane repair response, whose precise nature remains to be determined.

In conclusion, in this work we propose that host damage during *C. albicans* epithelial infection is driven by a ‘sequential’ damage mechanism in which candidalysin secretion acts to weaken host membranes enveloping invading hyphae prior to rupture, followed by hyphal traversal of two distinct subcellular niches, the JNR and the CCB, where membrane rupture occurs. As the results reported in this study are based on a highly simplified *in vitro* monolayer infection model using an immortalized epithelial cell line, their relevance to real-life *C. albicans* epithelial infection remains to be determined. Future research employing advanced *in vitro* as well as *in vivo* infection models should enable addressing the limitations imposed by the experimental model used here, while also testing the novel hypotheses and mechanisms arising from this study.

## Methods

### *C. albicans* strains and growth conditions

*C. albicans* BWP17+CIp30 (prototroph wildtype) (genotype rps1::(HIS1 ARG4 URA3) URA3::imm434/URA3::imm434 IRO1::imm434/IRO1::imm434 HIS1::hisG/HIS1::hisG chr5Part::Δ ARG4::hisG/ARG4::hisG RPS1::(HIS1 ARG4 URA3)) or ece1ha+CIp10 ece1 double knockout mutant (Genotype ece1::HIS1/ece1::ARG4 rps1::URA3 URA3::imm434/URA3::imm434 IRO1::imm434/IRO1::imm434 HIS1::hisG/HIS1::hisG chr5Part::Δ ARG4::hisG/ARG4::hisG ECE1::HIS1/ECE1::ARG4 RPS1::URA3)[6], kindly provided by Bernhard Hube (Leibniz Institute for Natural Product Research and Infection Biology-Hans Knoell Institute, Jena, Germany) were used as indicated. Strains were routinely grown on Yeast Extract Peptone Dextrose (YEPD) liquid/agar (1% yeast extract, 2% peptone, 2% D-glucose with or without 2% agar) supplemented with 80 μg/mL of uridine at 30°C.

### Epithelial cell lines culture

TR146 cell line expressing mOrange-Galectin-3 (Gal-3) was generated by Patricia Latour Lambert (Pasteur Institute, Paris, France) using the Sleeping Beauty plasmid (pSBBi-PUR[39]) with the coding sequences from the pEmOrangeGalectin3. Cells were routinely cultured in Dulbecco’s modified Eagle’s medium (DMEM) high glucose (Gibco-Invitrogen) supplemented with 10% fetal bovine calf serum (Gibco-Invitrogen) and 1% penicillin-streptomycin ((10,000 U/mL) Gibco) at 37°C, 5% CO2. Live-cell imaging was performed in optically transparent EM medium (120 mM NaCl, 7 mM KCl, 1.8 mM CaCl2, 0.8 mM MgCl2, 5 mM glucose and 25 mM Hepes at pH 7.3) supplemented with 0.2 g/L of amino acids (MP Biomedicals SC Amino Acids Mix) and 80 μg/mL of uridine[5].

### Infection procedure

For all experiments, TR146 cells were plated in a μ-Plate 96 Well Black (Ibidi 89626). 3×10^5^ cells/mL were plated 2 days before infection and were infected at 100% confluency. To label the host cell plasma membrane and endocytic compartment, CellMask Deep Red Plasma Membrane Stain (CM) (ThermoFisher) was used at a concentration of 2.5 μg/mL in EM media. TR146 cells were incubated with CM in the dark at 37°C for 10 minutes and then washed once with PBS (Gibco). The cell death marker SYTOX Green Nucleic Acid Stain (Invitrogen) was added at a concentration of 100 nM to EM media for the duration of the microscopy acquisition without washes. For infection, *C. albicans* strains were cultured overnight in liquid YEPD supplemented with 80 μg/mL uridine at 30°C, 200 rpm shaking, followed by a 1:100 back dilution into fresh YEPD and grown to exponential phase at 30°C, 200 rpm shaking for 4 hours. To prevent yeast from forming large clumps, the culture was subjected to mild sonication on ice for 5 cycles of 15 seconds on and 30 seconds off, at power setting 15% using a FisherBrand Model 50 Sonic Dismembrator (Fisher Scientific) with a standard ⅛” diameter microtip. Sonicated yeast cells were counted with a LUNA-FL reader (Logos Biosystems) and then diluted in EM medium and added to epithelial cells at a multiplicity of infection (MOI) 0.001 (1×10^3^ yeast cells/mL, 100 µL in each well). 30 min of ‘landing time’ to allow initial adherence of yeast to epithelial cells was followed by inoculating yeast (IY) identification, centering within the FOV and live cell imaging.

### Live-cell imaging

Live cell imaging was performed in a fully automated Inverted Nikon Ti2 Crest V3 spinning disk confocal equipped with Perfect focus system (PFS) and environmental chamber set to 37°C and 5% CO_2_. NIS elements (Nikon) was used for all data acquisition. For 24 hours acquisitions a 20X objective (NA 0.8) was used, with acquisition intervals of 30 min. For 12 hour acquisitions a 40X water immersion objective (NA 1.15) linked to a water dispenser (Nikon) was used, with acquisition intervals of 10 min. Z stack of 12 planes, 1 μm step was used, with 8 FOVs acquired per condition (WT, *ece1*Δ/Δ) plus 4 FOVs for non-infected controls in 24-hour acquisitions. For the Bright-field (BF) channel, only a single central z-section was acquired. For “high-resolution” 2-hour acquisitions, a 60X oil immersion objective (NA 1.42) was used, with acquisition intervals of 2 min. Z stack of 13 planes, 0.5 μm step was used, with 6 FOVs acquired of WT infection. Imaging was started 5 hours post-infection for 2 hours total. 20X objective acquisitions were used for analysis in Figure 1D. 40X objective acquisitions are presented and used for analysis in Figure 1C, 3A, 3B, 3C, 4C, 4D, 4E. 60X objective acquisitions are presented in Figures 4A, 4B, 5A, 5B. Supplementary movies correspond to the objectives used in figures.

### Optical z-sectioning at high resolution

TR146 invaded by hyphae were first examined live (prior to fixation) using the CM channel at around 7 hours post-infection. A 100X oil immersion objective (NA 1.45) was used to acquire z-stacks of around 100 slices with a z-step of 0.2 μm. The positions of acquired infection sites were saved in the microscope software (NIS-Elements, Nikon), at which point the sample was fixed using PFA 4% for 10 min at room temperature. The sample was then stained with calcofluor white (Fluorescent brightener 28, MP Biomedicals) at a concentration of 5 µg/ml for 10 min, followed by a single wash in PBS. Finally, the sample was stained with Phalloidin-FITC (Sigma), at a concentration of 0.1 µg/ml for 1 hour, followed by three washes in PBS. The acquired positions were then re-identified in the microscope, followed by identical z-stack acquisitions in the blue (CFW), green (Phalloidin-FITC) and far-red (CM) channels. 100X acquisitions are presented in Fig 2A, 2B, 2C, S3 Fig.

### Serial-block face scanning electron microscopy

For Serial Block Face-Scanning Electron Microscopy (SBF-SEM), TR146 cell line expressing mOrange-Gal-3 were plated to confluency on 35 mm cell culture treated (ibiTreat) plastic bottom dishes (Ibidi 81156) and infected with *C. albicans* BWP17+CIp30 or ece1ha+CIp10 ece1 double knockout mutant for 5 h at MOI 0.1, then fixed in 3% PFA, 1% glutaraldehyde during 1 h at room temperature. To increase conductivity of the sample in the microscope, the cell monolayer was covered with a thin layer of 10% gelatine containing 2% of Bovine Serum Albumine (BSA). Samples were then prepared for SBF-SEM (NCMIR protocol[40]) as follows: cells were post-fixed for 1 h in a reduced osmium solution containing 2% osmium tetroxide, 1.5% potassium ferrocyanide in PBS, followed by incubation with a 1% thiocarbohydrazide in water for 20 min. Subsequently, samples were stained with 2% OsO4 in water for 30 min, followed by 1% aqueous uranyl acetate at 4 °C overnight. Cell monolayers were then subjected to en bloc Walton’s lead aspartate staining[41], and placed in a 60 °C oven for 30 min. Samples were then dehydrated in graded concentrations of ethanol for 15 min in each step. The samples were infiltrated with 30% agar low viscosity resin (Agar Scientific Ltd, UK) in ethanol, for 1 h, 50% resin for 2 h and 100% resin overnight. The resin was then changed and the cells were further incubated during 5.5 h, prior to inclusion in upside down capsules and polymerization for 18 h at 60 °C. The polymerized blocks were mounted onto aluminum stubs together with plastic substrate for SBF imaging (FEI Microtome 8 mm SEM Stub, Agar Scientific), with two-part conduction silver epoxy kit (EMS, 12642-14). For imaging, samples on aluminum stubs were trimmed using an ultramicrotome and inserted into a TeneoVS SEM (ThermoFisher). Acquisitions were performed with a beam energy of 2.7 kV, 400 pA current, in LowVac mode at 40 Pa, a dwell time of 2 µs per pixel at 10 nm pixel size. Sections of 100 nm were serially cut between images. Overall, 2 datasets (WT and *ece1*Δ/Δ) were acquired, with 5 different fields of view with a total of 1749 slices for WT, and 5 different fields of view with a total of 2293 slices for the *ece1*Δ/Δ.

### Image processing and analysis

Light microscopy data was analyzed using NIS-Elements (Nikon), Fiji[42] and CellProfiler[43]. Three-dimensional visualization of hyphal invasion (Figure 2C) was generated using NIS-Elements. Galectin-3 recruitment sites were identified and annotated manually using NIS-Elements. Host cell death as indicated by SYTOX Green was quantified using CellProfiler using a custom pipeline described in **S1 Text**. SBF-SEM data was processed using Fiji and Amira (ThermoFisher). Data alignment and manual segmentation were performed using Amira. Figures were prepared using NIS-Elements, Fiji and Inkscape.

### Quantifications, statistical analysis and reproducibility

24 hours live cell imaging data were acquired in three independent experiments. 12 hours live cell imaging data were acquired in four independent experiments. For all quantifications of Gal-3 recruitment, only the first 10 hours of each acquisition were analysed. 2 hours live cell imaging data was acquired in two independent experiments. Data was analysed using RStudio. p-value thresholds: 0.05 (*), 0.01 (**), 0.001 (***). χ2 test for homogeneity was performed in Figure 3B, 4C, 4D, and 4E. Student t-test was performed in Figure 3C, error bars represent SEM. The numerical raw data used to generate quantifications in all figures is available in **S1 Appendix**.

## Supporting information

S1 Appendix. Raw numerical data used for generating quantifications

## Data availability

All data from this study are available from the corresponding author upon request.

## Acknowledgements

We thank Bernhard Hube (Leibniz Institute for Natural Product Research and Infection Biology-Hans Knoell Institute, Jena, Germany) for providing us with the *C. albicans* strains used in this study. We thank Jost Enninga (Pasteur Institute, Paris, France), for support in production of stably expressing epithelial cell lines. We thank Robert Arkowitz (IBV, Université Côte d’Azur, Nice France) for critical reading of the manuscript. We thank UMS28 in Sorbonne University for support with BSL2 facility and microscopy. We acknowledge the ImagoSeine core facility of the Institut Jacques Monod, member of the France BioImaging infrastructure supported by the French National Research Agency (ANR-24-INBS-0005 FBI BIOGEN) and GIS-IBiSA, and the support of the Région Île-de-France (Sesame). This work was supported by the ATIP-Avenir program to A.W., and by the Région Île-de-France via the DIM1Health action for funding of the spinning disk confocal microscope used in this study and PhD funding to L.M.

## Contributions

Conceptualization, A.W.; Methodology, L.M., N.C., A.S., P.L., J-F.F, C.P-D., R.L.B, J-M.V, M.L. and .A W.; Investigation, L.M., N.C., A.S., C.P-D., R.L.B, and A.W.; Formal Analysis, N.C., L.M., M.L. and A.W.; Writing - Original Draft, A.W.; Supervision, A.W.

## Supporting movie captions

**S1 Movie. Corresponding to Fig 1C**. Host membrane rupture as indicated by galectin-3 recruitments during the first 12 hours of WT TR146 infection. From left to right: Bright-field (BF), SYTOX green (green), Gal-3 (red) and CellMask (CM) (magenta). Individual Gal-3 recruitments are highlighted with boxes in the Gal-3 channel. Scale bar is 100 µm.

**S2 Movie. Corresponding to Fig 1D**. Host cell death as indicated by SYTOX green staining during 24 hours of WT TR146 infection. From left to right: Bright-field (BF), SYTOX green (green), Gal-3 (red) and CellMask (CM) (magenta). Scale bar is 100 µm.

**S3 Movie. Corresponding to Fig 2A**. High-resolution z-stack acquired with 100X oil immersion objective of a (live) non-fixed invasion site labelled with CellMask. Z interval is 0.2 µm. Scale bar is 20 µm.

**S4 movie. Corresponding to Fig 2B**. High-resolution z-stack acquired with 100X oil immersion objective of a fixed invasion site labelled with (from left to right) CFW (fungal cell wall), Phalloidin (Actin) and CellMask. A composite view is also provided (far right). Z interval is 0.2 µm. Scale bar is 20 µm.

**S5 movie. Corresponding to Fig 2D**. Serial block face-scanning electron microscopy (SBF-SEM) of WT invasion site. An XY section through the volume is presented, followed by segmentation of a hypha (white), the nucleus (teal) and the epithelial cell membrane (yellow).

**S6 movie. Corresponding to Fig 2E**. Serial block face-scanning electron microscopy (SBF-SEM) of a WT hypha invading two cells in sequence. Intra- and inter-cellular invasion is observed as well as ‘complete’ and ‘partial’ invasion pocket geometries. White - hypha, yellow - Cell 1, teal - Cell 1 nucleus, magenta-Cell 2.

**S7 movie, corresponding to Fig 3A**. ‘Tip’ Gal-3 recruitment pattern. From left to right: Gal-3, CellMask (CM) and merge of Gal-3 (red) and CM (white) channels. Scale bar is 10 µm.

**S8 movie, corresponding to Fig 3A**. ‘Tubular’ Gal-3 recruitment pattern. From left to right: Gal-3, CellMask (CM) and merge of Gal-3 (red) and CM (white) channels. Scale bar is 10 µm.

**S9 movie, corresponding to Fig 3A**. ‘Elongated’ Gal-3 recruitment pattern. From left to right: Gal-3, CellMask (CM) and merge of Gal-3 (red) and CM (white) channels. Scale bar is 10 µm.

**S10 movie, corresponding to Fig 4**. Long hyphal trajectory through two host cells, with both JNR and CCB Gal-3 recruitments observed in sequence. From left to right: Gal-3, CellMask (CM) and merge of Gal-3 (red) and CM (white) channels. Scale bar is 10 µm. The timeline corresponds to that of the associated main figure. Imaging intervals are 2 min.

**S11 movie, corresponding to Fig 5A. -** CCB rupture captured with high-spatial and -temporal resolution using 60X oil immersion objective. From left to right: Gal-3, CellMask (CM) and merge of Gal-3 (red) and CM (white) channels. Scale bar is 10 µm. The timeline corresponds to that of the associated main figure. Imaging intervals are 2 min.

**S12 movie, corresponding to Fig 5B. -** JNR rupture captured with high-spatial and -temporal resolution using 60X oil immersion objective. From left to right: Gal-3, CellMask (CM) and merge of Gal-3 (red) and CM (white) channels. Scale bar is 10 µm. The timeline corresponds to that of the associated main figure. Imaging intervals are 2 min.

## References

1. Gow NAR, van de Veerdonk FL, Brown AJP, Netea MG. Candida albicans morphogenesis and host defence: discriminating invasion from colonization. Nat Rev Microbiol. 2011. doi:10.1038/nrmicro2711

2. Brown GD, Denning DW, Gow NAR, Levitz SM, Netea MG, White TC. Hidden killers: human fungal infections. Sci Transl Med. 2012;4. doi:10.1126/SCITRANSLMED.3004404

3. Höfs S, Mogavero S, Hube B. Interaction of Candida albicans with host cells: virulence factors, host defense, escape strategies, and the microbiota. Journal of Microbiology. The Microbiological Society of Korea; 2016. pp. 149–169. doi:10.1007/s12275-016-5514-0

4. Dalle F, Wächtler B, L’Ollivier C, Holland G, Bannert N, Wilson D, et al. Cellular interactions of Candida albicans with human oral epithelial cells and enterocytes. Cell Microbiol. 2010;12: 248–271. doi:10.1111/j.1462-5822.2009.01394.x

5. Lachat J, Pascault A, Thibaut D, Le Borgne R, Verbavatz J-M, Weiner A. Trans-cellular tunnels induced by the fungal pathogen Candida albicans facilitate invasion through successive epithelial cells without host damage. Nat Commun. 2022;13: 3781. doi:10.1038/s41467-022-31237-z

6. Moyes DL, Wilson D, Richardson JP, Mogavero S, Tang SX, Wernecke J, et al. Candidalysin is a fungal peptide toxin critical for mucosal infection. Nature. 2016;532: 64–68. doi:10.1038/nature17625

7. Mogavero S, Sauer FM, Brunke S, Allert S, Schulz D, Wisgott S, et al. Candidalysin delivery to the invasion pocket is critical for host epithelial damage induced by Candida albicans. Cell Microbiol. 2021;23. doi:10.1111/cmi.13378

8. Wilson D, Naglik JR, Hube B. The Missing Link between Candida albicans Hyphal Morphogenesis and Host Cell Damage. Hogan DA, editor. PLoS Pathogens. Public Library of Science; 2016. p. e1005867. doi:10.1371/journal.ppat.1005867

9. Lin J, Miao J, Schaefer KG, Russell CM, Pyron RJ, Zhang F, et al. Sulfated glycosaminoglycans are host epithelial cell targets of the Candida albicans toxin candidalysin. Nature Microbiology 2024 9:10. 2024;9: 2553–2569. doi:10.1038/s41564-024-01794-8

10. Russell CM, Schaefer KG, Dixson A, Gray AL, Pyron RJ, Alves DS, et al. The Candida albicans virulence factor candidalysin polymerizes in solution to form membrane pores and damage epithelial cells. Elife. 2022;11. doi:10.7554/elife.75490

11. Allert S, Förster TM, Svensson CM, Richardson JP, Pawlik T, Hebecker B, et al. Candida albicans-induced epithelial damage mediates translocation through intestinal barriers. mBio. 2018;9. doi:10.1128/mBio.00915-18

12. Richardson JP, Moyes DL, Ho J, Naglik JR. Candida innate immunity at the mucosa. Semin Cell Dev Biol. 2018 [cited 28 Sep 2018]. doi:10.1016/j.semcdb.2018.02.026

13. Wächtler B, Citiulo F, Jablonowski N, Förster S, Dalle F, Schaller M, et al. Candida albicans-epithelial interactions: dissecting the roles of active penetration, induced endocytosis and host factors on the infection process. PLoS One. 2012;7: e36952. doi:10.1371/journal.pone.0036952

14. Westman J, Plumb J, Licht A, Yang M, Allert S, Naglik JR, et al. Calcium-dependent ESCRT recruitment and lysosome exocytosis maintain epithelial integrity during Candida albicans invasion. Cell Rep. 2022;38. doi:10.1016/j.celrep.2021.110187

15. Almeida RS, Brunke S, Albrecht A, Thewes S, Laue M, Edwards JE, et al. the hyphal-associated adhesin and invasin Als3 of Candida albicans mediates iron acquisition from host ferritin. PLoS Pathog. 2008;4: e1000217. doi:10.1371/journal.ppat.1000217

16. Paz I, Sachse M, Dupont N, Mounier J, Cederfur C, Enninga J, et al. Galectin-3, a marker for vacuole lysis by invasive pathogens. Cell Microbiol. 2010;12: 530–544. doi:10.1111/j.1462-5822.2009.01415.x

17. Weiner A, Mellouk N, Lopez-Montero N, Chang Y-Y, Souque C, Schmitt C, et al. Macropinosomes are Key Players in Early Shigella Invasion and Vacuolar Escape in Epithelial Cells. PLoS Pathog. 2016;12: e1005602. doi:10.1371/journal.ppat.1005602

18. Ellison CJ, Kukulski W, Boyle KB, Munro S, Randow F. Transbilayer Movement of Sphingomyelin Precedes Catastrophic Breakage of Enterobacteria-Containing Vacuoles. Current Biology. 2020;30: 2974–2983.e6. doi:10.1016/j.cub.2020.05.083

19. Gutierrez MG, Enninga J. Intracellular niche switching as host subversion strategy of bacterial pathogens. Current Opinion in Cell Biology. Elsevier Ltd; 2022. doi:10.1016/j.ceb.2022.102081

20. Richardson JP, Willems HME, Moyes DL, Shoaie S, Barker KS, Tan SL, et al. Candidalysin Drives Epithelial Signaling, Neutrophil Recruitment, and Immunopathology at the Vaginal Mucosa. Infect Immun. 2018;86. doi:10.1128/IAI.00645-17

21. Watts HJ, Véry AA, Perera THS, Davies JM, Gow NAR. Thigmotropism and stretch-activated channels in the pathogenic fungus Candida albicans. Microbiology (Reading). 1998;144 ( Pt 3): 689–695. doi:10.1099/00221287-144-3-689

22. Brand A, Vacharaksa A, Bendel C, Norton J, Haynes P, Henry-Stanley M, et al. An internal polarity landmark is important for externally induced hyphal behaviors in Candida albicans. Eukaryot Cell. 2008;7: 712–20. doi:10.1128/EC.00453-07

23. Thomson DD, Wehmeier S, Byfield FJ, Janmey PA, Caballero-Lima D, Crossley A, et al. Contact-induced apical asymmetry drives the thigmotropic responses of *C andida albicans* hyphae. Cell Microbiol. 2015;17: 342–354. doi:10.1111/cmi.12369

24. Schaller M, Zakikhany K, Naglik JR, Weindl G, Hube B. Models of oral and vaginal candidiasis based on in vitro reconstituted human epithelia. Nat Protoc. 2006;1: 2767–73. doi:10.1038/nprot.2006.474

25. Russell CM, Rybak JA, Miao J, Peters BM, Barrera FN. Candidalysin: Connecting the pore forming mechanism of this virulence factor to its immunostimulatory properties. J Biol Chem. 2023;299. doi:10.1016/J.JBC.2022.102829

26. Fu R, Jiang X, Li G, Zhu Y, Zhang H. Junctional complexes in epithelial cells: sentinels for extracellular insults and intracellular homeostasis. FEBS J. 2022;289: 7314–7333. doi:10.1111/FEBS.16174

27. Puerner C, Kukhaleishvili N, Thomson D, Schaub S, Noblin X, Seminara A, et al. Mechanical force-induced morphology changes in a human fungal pathogen. BMC Biol. 2020;18. doi:10.1186/S12915-020-00833-0

28. Lenne PF, Rupprecht JF, Viasnoff V. Cell Junction Mechanics beyond the Bounds of Adhesion and Tension. Dev Cell. 2021;56: 202–212. doi:10.1016/J.DEVCEL.2020.12.018

29. Phan QT, Myers CL, Fu Y, Sheppard DC, Yeaman MR, Welch WH, et al. Als3 is a Candida albicans invasin that binds to cadherins and induces endocytosis by host cells. PLoS Biol. 2007;5: e64. doi:10.1371/journal.pbio.0050064

30. Sun JN, Solis N V., Phan QT, Bajwa JS, Kashleva H, Thompson A, et al. Host Cell Invasion and Virulence Mediated by Candida albicans Ssa1. Levitz SM, editor. PLoS Pathog. 2010;6: e1001181. doi:10.1371/journal.ppat.1001181

31. Villar CC, Kashleva H, Nobile CJ, Mitchell AP, Dongari-Bagtzoglou A. Mucosal tissue invasion by Candida albicans is associated with E-cadherin degradation, mediated by transcription factor Rim101p and protease Sap5p. Infect Immun. 2007;75: 2126–2135. doi:10.1128/IAI.00054-07

32. Frank CF, Hostetter MK. Cleavage of E-cadherin: a mechanism for disruption of the intestinal epithelial barrier by Candida albicans. Transl Res. 2007;149: 211–222. doi:10.1016/J.TRSL.2006.11.006

33. Goyer M, Loiselet A, Bon F, L’Ollivier C, Laue M, Holland G, et al. Intestinal Cell Tight Junctions Limit Invasion of Candida albicans through Active Penetration and Endocytosis in the Early Stages of the Interaction of the Fungus with the Intestinal Barrier. PLoS One. 2016;11: e0149159. doi:10.1371/journal.pone.0149159

34. Böhringer M, Pohlers S, Schulze S, Albrecht-Eckardt D, Piegsa J, Weber M, et al. Candida albicans infection leads to barrier breakdown and a MAPK/NF-κB mediated stress response in the intestinal epithelial cell line C2BBe1. Cell Microbiol. 2016;18: 889–904. doi:10.1111/cmi.12566

35. Sprague JL, Schille TB, Allert S, Trümper V, Lier A, Großmann P, et al. Candida albicans translocation through the intestinal epithelial barrier is promoted by fungal zinc acquisition and limited by NFκB-mediated barrier protection. PLoS Pathog. 2024;20. doi:10.1371/JOURNAL.PPAT.1012031

36. Lapaquette P, Ducreux A, Basmaciyan L, Paradis T, Bon F, Bataille A, et al. Membrane protective role of autophagic machinery during infection of epithelial cells by *Candida albicans*. Gut Microbes. 2022;14. doi:10.1080/19490976.2021.2004798

37. Wayakanon K, Thornhill MH, Douglas CWI, Lewis AL, Warren NJ, Pinnock A, et al. Polymersome-mediated intracellular delivery of antibiotics to treat Porphyromonas gingivalis-infected oral epithelial cells. FASEB J. 2013;27: 4455–4465. doi:10.1096/FJ.12-225219

38. Biwer LA, Isakson BE. Endoplasmic reticulum mediated signaling in cellular microdomains. Acta Physiol (Oxf). 2016;219: 162. doi:10.1111/APHA.12675

39. Kowarz E, Löscher D, Marschalek R. Optimized Sleeping Beauty transposons rapidly generate stable transgenic cell lines. Biotechnol J. 2015;10: 647–653. doi:10.1002/BIOT.201400821

40. Deerinck TJ, Bushong E a., Thor A, Ellisman MH. NCMIR methods for 3D EM: A new protocol for preparation of biological specimens for serial block face scanning electron microscopy. Microscopy. 2010; 6–8.

41. Walton J. Lead aspartate, an en bloc contrast stain particularly useful for ultrastructural enzymology. Journal of Histochemistry and Cytochemistry. 1979;27: 1337–1342. doi:10.1177/27.10.512319

42. Schindelin J, Arganda-Carreras I, Frise E, Kaynig V, Longair M, Pietzsch T, et al. Fiji: An open-source platform for biological-image analysis. Nature Methods. Nat Methods; 2012. pp. 676–682. doi:10.1038/nmeth.2019

43. Stirling DR, Swain-Bowden MJ, Lucas AM, Carpenter AE, Cimini BA, Goodman A. CellProfiler 4: improvements in speed, utility and usability. BMC Bioinformatics. 2021;22. doi:10.1186/S12859-021-04344-9

