## Supplementary information and figures for "Membrane damage during *Candida albicans* epithelial invasion is localized to distinct host subcellular niches"

**S1 Text. Supporting information materials and methods**

**CellProfiler pipeline used for host cell death quantification using SYTOX green (Fig 1D)**

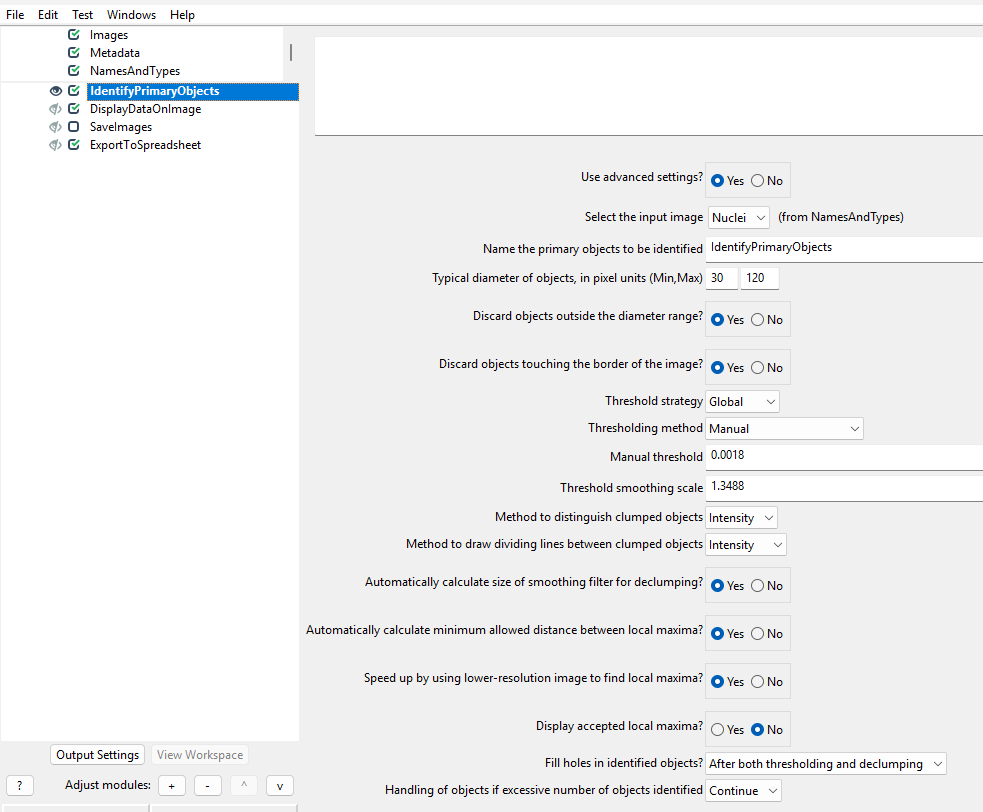

**Supplementary Figures S1, S2, S3, S4**

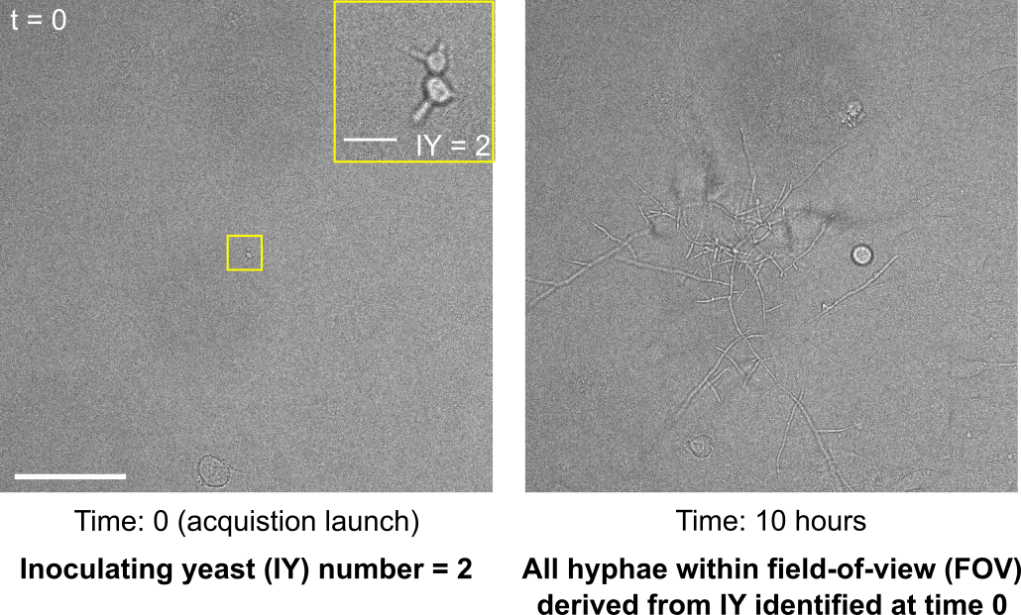

**S1 Fig. Tracking of inoculating yeasts (IYs) over time within a distinct ‘infection site’.** At the start of infection (t = 0), sites containing between one and four inoculating yeasts (IYs) are identified and centred to the middle of the microscope field of view (FOV), followed by live cell imaging of multiple FOVs in parallel. All subsequent infection events (e.g. membrane rupture, host cell death) observed are derived from the identified IY at time 0, comprising a single ‘infection site’. Images acquired using 40X objective.

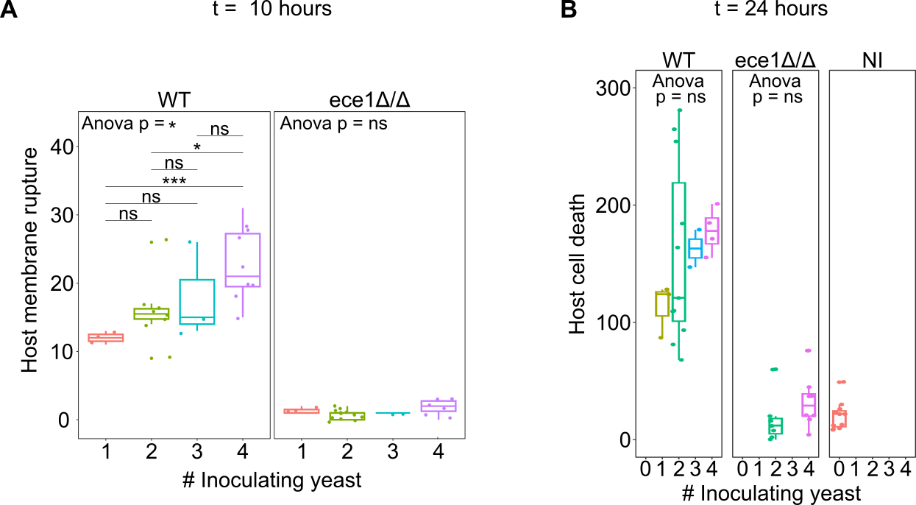

**S2 Fig. Correlation between fungal load and host membrane rupture or host cell death during WT infection**. (A) Total host membrane rupture as indicated by Gal-3 recruitments at 10 hours of infection plotted against the inoculating yeast (IY) number. (B) Total host cell deaths as indicated by SYTOX green labeling events at 24 hours of infection plotted against the inoculating yeast number. ANOVA test on all groups and student t-test are presented, p-value thresholds: 0.05 (*), 0.01 (**), 0.001 (***).

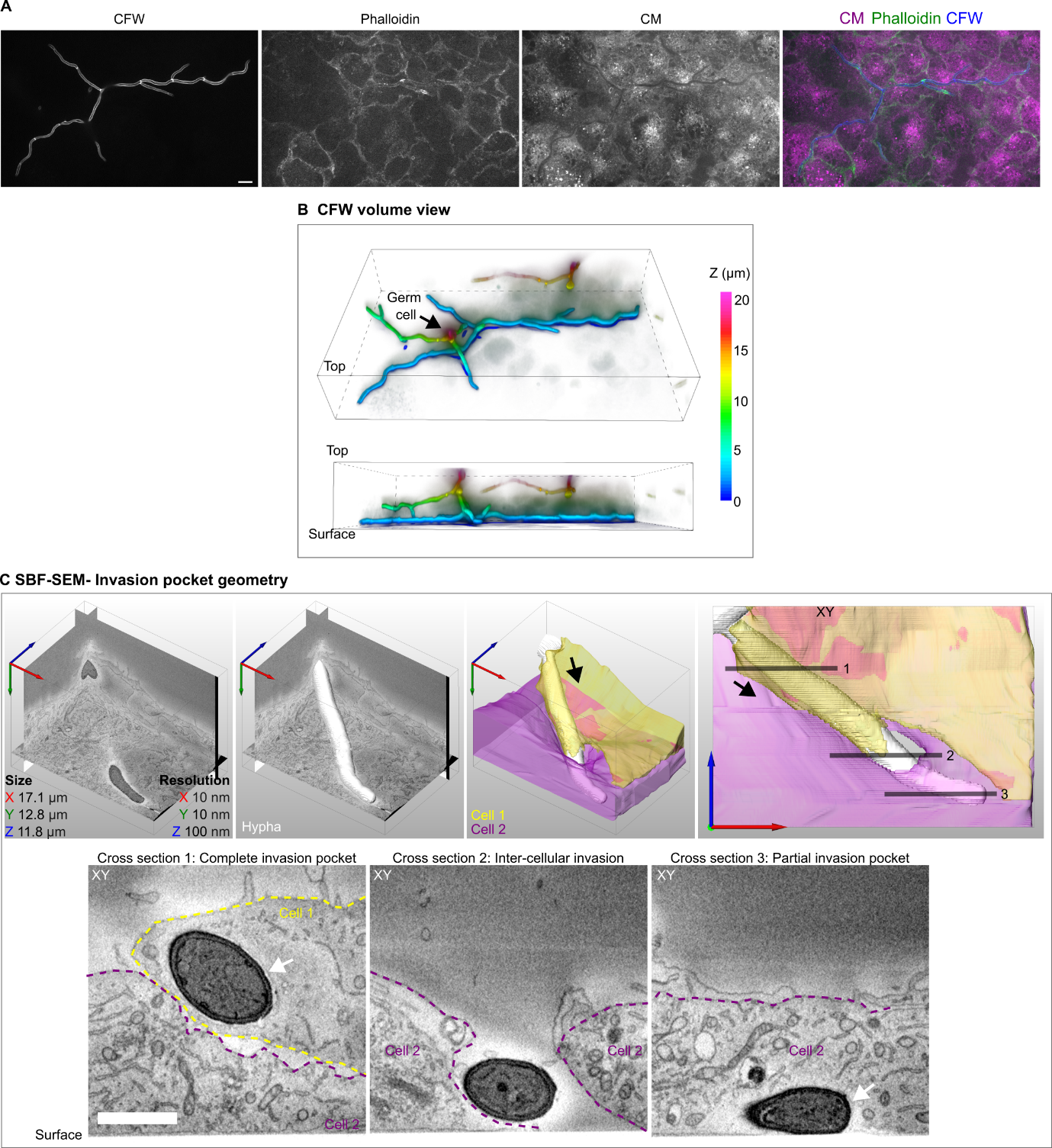

**S3 Fig. Invasion architecture of ece1Δ/Δ strain.** (A) Fixed ece1Δ/Δ invasion site stained with CFW, a fungal cell wall marker, phalloidin, an actin marker and CM. In all images the optical section nearest to the microplate surface is shown. (B) A three-dimensional representation of CFW staining in panel B using z-stack color coding. (C) SBF-SEM dataset of an *ece1*Δ/Δ hypha (white) invading two host cells (Cell 1- yellow, Cell 2 - magenta) in sequence. XY cross-sections from within the volume are presented at three different planes along the invasion trajectory: cross section 1 - a ‘complete’ invasion pocket surrounding the hypha segment within Cell 1; cross section 2 - a hypha segment within the inter-cellular space; cross section 3 - a ‘partial’ invasion pocket around a hypha segment in direct contact with the microplate surface within Cell 2. The invasion pocket membrane is indicated with a white arrow. In all panels, the direction of hyphal extension is indicated with black arrows. Scale bars are 10 µm in A and 2 µm in C. The dimensions and resolution of the SBF-SEM volume presented is listed within the figure.

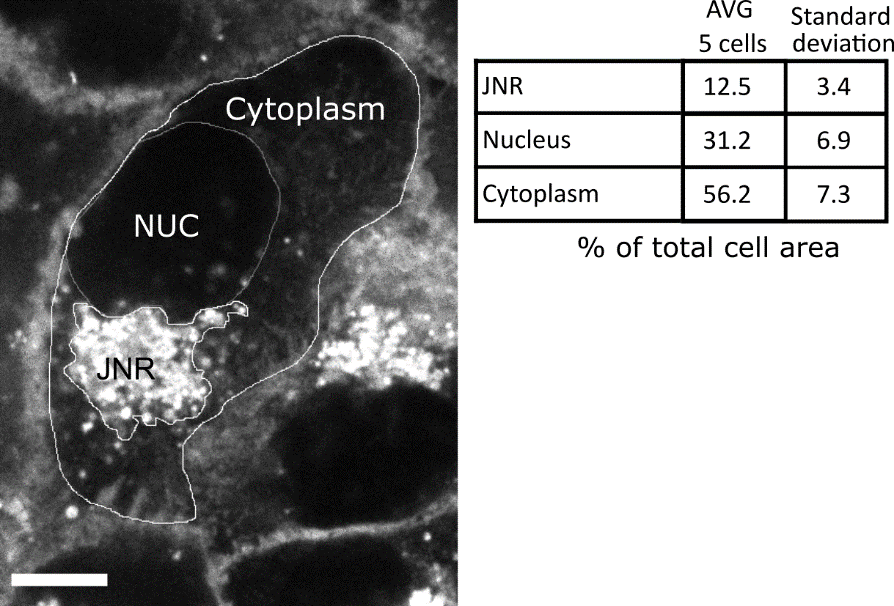

**S4 Fig. Estimation of dimensions of different compartments within TR146 cells.** A representative z section of a cell labelled with CellMask is presented, with the juxtanuclear region (JNR), the nucleus (NUC) and cytoplasm outlined. An average and standard deviation of the relative areas of each compartment in five different cells is presented. Scale bar is 10 µm. Analysis was performed using data acquired with a 60X oil immersion objective.
